# A rare *SMAD9* mutation identifies the BMP signalling pathway as a potential osteoanabolic target

**DOI:** 10.1101/560565

**Authors:** Celia L Gregson, Dylan Bergen, Paul Leo, Richard B. Sessions, Lawrie Wheeler, April Hartley, Scott Youlten, Peter I Croucher, Aideen M. McInerney-Leo, William Fraser, Jonathan C.Y. Tang, Lisa Anderson, Mhairi Marshall, Leon Sergot, Lavinia Paternoster, George Davey-Smith, The AOGC Consortium, Matthew A Brown, Chrissy Hammond, John P Kemp, Jon H Tobias, Emma L Duncan

**Affiliations:** Musculoskeletal Research Unit, Translational Health Sciences, Bristol Medical School, University of Bristol, Bristol, UK; School of Physiology, Pharmacology, and Neuroscience, Faculty of Life Sciences, University of Bristol, UK; Translational Genomics Group, Institute of Health and Biomedical Innovation, Faculty of Health, Queensland University of Technology, Translational Research Institute, Princess Alexandra Hospital, Ipswich Rd, Woolloongabba, QLD 4102, Australia.; School of Biochemistry, Faculty of Life Sciences, University of Bristol, UK; Translational Genomics Group, Institute of Health and Biomedical Innovation, School of Biomedical Sciences, Queensland University of Technology (QUT), Translational Research Institute, 37 Kent St, Woolloongabba, QLD, 4102.; Medical Research Council Integrative Epidemiology Unit, Population Health Sciences, Bristol Medical School, University of Bristol, UK; Division of Bone Biology, Garvan Institute of Medical Research, Sydney, NSW 2010, Australia; St Vincent’s Clinical School, Faculty of Medicine, UNSW Sydney, Sydney NSW 2010, Australia; School of Biotechnology and Biomolecular Sciences, UNSW Sydney, Sydney, NSW 2052, Australia; Dermatology Research Centre, The University of Queensland, The University of Queensland Diamantina Institute, Brisbane, Queensland, Australia.; Norwich Medical School, University of East Anglia, Norwich Research Park, Norwich, UK.; Departments of Diabetes, Endocrinology and Clinical Biochemistry, Norfolk and Norwich University Hospital NHS Foundation Trust, Colney Lane, Norwich, UK; Severn School of Radiology, Severn Deanery, Park House, Bristol, UK; The University of Queensland Diamantina Institute, Translational Research Institute, University of Queensland, Brisbane, QLD 4102, Australia; Department of Endocrinology and Diabetes, Royal Brisbane & Women’s Hospital, Butterfield St, Herston, QLD 4029, Australia.; Faculty of Medicine, University of Queensland, Herston, QLD 4006, Australia.

**Keywords:** High Bone Mass, SMAD9, DXA, exon sequencing, monogenic, zebrafish

## Abstract

To identify targets for novel anabolic medicines for osteoporosis, we recruited a large cohort with unexplained high bone mass (HBM). Exome sequencing identified a rare (minor allele frequency 0.0014) missense mutation in *SMAD9* (c.65T>C, p.Leu22Pro) segregating with HBM in an autosomal dominant family. The same mutation was identified in another two unrelated individuals with HBM. *In-silico* protein modelling predicts the mutation severely disrupts the MH1 DNA-binding domain of SMAD9. Affected individuals have bone mineral density [BMD] Z-Scores +3 to +5, with increased volumetric cortical and trabecular BMD, increased cortical thickness, and low/normal bone turnover. Fractures and nerve compressions are not seen. Both genome-wide, and gene-based association testing of heel estimated-BMD in >362,924 UK-Biobank British subjects showed strong associations with *SMAD9* (P_GWAS_=6×10^−16^; P_GENE_ =8×10^−17^). Smad9 is highly expressed in murine osteocytes and zebrafish bone tissue. Our findings support *SMAD9* as a novel HBM gene, and a potential novel osteoanabolic target.

## Background

Age-related bone loss with deterioration of skeletal architecture leads to osteoporosis, affecting 8.2 million women and 2.0 million men aged 50 years and older in the United States (US) ^1^. Worldwide, osteoporosis causes more than 8.9 million fractures annually ^1^. Osteoporotic fractures and their treatment are a major cause of morbidity and mortality, with annual US healthcare costs exceeding $20 billion ^2^. Most osteoporosis treatment approaches, including all oral medications, reduce bone resorption and prevent further bone loss, rather than enhance bone formation. Affordable anabolic treatments, which can restore bone mass and skeletal architecture, are much needed.

Romosozumab, a monoclonal antibody against sclerostin, represents a new class of anti-osteoporosis drugs ^3,4^ (licensing currently under review). Sclerostin, a key inhibitor of bone formation, was discovered through study of two rare syndromes of extreme high bone mass (HBM) due to mutations in *SOST* ^5,6^. *SOST* encodes Sclerostin which binds to low-density lipoprotein receptor-related proteins 5 and 6 (LRP5 and LRP6) to prevent activation of canonical WNT signalling in bone, resulting in decreased bone formation. Gain-of-function mutations in *LRP5* and *LRP6* can also cause extreme HBM ^7,8^. Together these sclerosing bone dysplasias are characterised by mandible enlargement with tori of the palate and mandible, bone overgrowth leading to nerve compression, a tendency to sink when swimming, and, importantly, resistance to fracture ^5,7,9^. These important gene discoveries validate the study of rare monogenic HBM as an approach to identify novel therapeutic targets for drug development towards osteoporosis treatments.

We have previously shown that HBM (defined as a total hip and/or 1^st^ lumbar vertebral bone mineral density [BMD] Z-score of ≥ +3.2) is observed in 0.18% of Dual-Energy X-ray Absorptiometry (DXA) scans in the UK ^10^. Most cases are unexplained, *i.e*. they do not carry mutations in established HBM genes ^9^. Whilst such HBM populations do show enrichment for common variant associations with established BMD-associated loci ^11^, we hypothesized that novel causes of monogenic HBM remain to be determined. Thus, we aimed to identify novel monogenic causes of HBM to provide insight into regulatory pathways amenable to therapeutic intervention.

## Results

### HBM pedigree with a segregating *SMAD9 p*.Leu22Pro mutation (Figure 1 and Table 1)

We investigated a pedigree with unexplained and apparently autosomal dominant HBM ^9^, identified from our large UK HBM cohort ^10^ (Figure 1).

**Table 1:** Characteristics of the c.65T>C, p.Leu22Pro *SMAD9* HBM pedigree members and two further unrelated HBM individuals with the same *SMAD9* mutation.

|  | HBM Pedigree |  |  |  | Additional Isolated HBM cases |  |
| --- | --- | --- | --- | --- | --- | --- |
|  | UK proband III.1 | UK half sister III.2 | UK mother II.2 | UK grand-mother I.2 (unaffected) | UK case | Australian case |
| ***SMAD9* Mutation** | Leu22Pro | Leu22Pro | Leu22Pro | WT | Leu22Pro | Leu22Pro |
| Age at assessment | 33 | 22 | 55 | 75 | 55 | 57 |
| Sex | Female | Female | Female | Female | Female | Female |
| Ethnicity | Caucasian | Caucasian | Caucasian | Caucasian | Caucasian | Caucasian |
| **Anthropometry** |  |  |  |  |  |  |
| Height (cm) | 178.0 | 173.3 | 175.0 | 160.6 | 160.0 | 162.2 |
| Weight (kg) | 138.0 | 133.8 | 127.8 | 72.6 | 89.7 | 69.9 |
| BMI (kg/m<sup>2</sup>) | 43.6 | 43.3 | 41.4 | 28.1 | 35.0 | 26.6 |
| **DXA Measurements** |  |  |  |  |  |  |
| Total Hip BMD Z-score | 3.2 | 4.8 | 3.3 | 0.1 | 4.3 | 2.7 |
| L1 BMD Z-score | 4.5 | 2.6 | 3.3 | 0.8 | 5.0 | +3.4* |
| BMC (kg) | 3.49 | 3.77 | 3.65 | 2.12 | 3.24 | - |
| Fat mass (kg) | 73.2 | 64.5 | 64.8 | 25.4 | 34.0 | - |
| Lean mass (kg) | 61.3 | 65.6 | 59.5 | 45 | 52.4 | - |
| **Clinical phenotype** |  |  |  |  |  |  |
| Adult Fracture | No | No | No | No | No | No |
| Sinks/ Floats | Floats | Sinks | Floats | Floats | Sinks | - |
| Bone pain | Yes | Yes | Yes | No | No | - |
| Visual/auditory impairment | Myopia | No | No | Impaired hearing | Astigmatism | - |
| Dentition | Normal | Normal | Normal | Normal | Retained cuspid tooth | - |
| Shoe Size <sup>a</sup> | 10 | 10 | 9 | 6 | 5.5 | - |
| Broad Frame | Yes | Yes | Yes | No | Yes | - |
| Enlarged Mandible | Yes | Yes | Yes | No | Yes | - |
| Torus | Yes | Yes | No | No | No | - |
| Nerve compression | No | No | No | No | No | - |
| Pes planus | No | No | Yes | No | Yes | - |
| **Blood tests** |  |  |  |  |  |  |
| ALP (IU/L) | 99 | 83 | 102 | 202 | 61 | - |
| Adjusted calcium (mmol/L) | 2.50 | 2.46 | 2.40 | 2.46 | 2.33 | - |
| P1NP (ug/L) | 58 | 36 | 22 | 95 | 35 | - |
| CTX (ug/L) | 0.15 | 0.18 | 0.19 | 0.10 | 0.16 | - |
| Osteocalcin (ug/L) | 12.4 | 17.1 | 13.5 | 14.8 | 11.5 | - |
| Sclerostin (pmol/L) | 71.0 | - | 56.1 | 44.4 | 50.4 | - |

WT: Wild type. <sup>a</sup> UK measurements. Reference ranges; ALP 20-120, Adjusted calcium 2.25-2.70, P1NP: pre-menopausal 30-78 ug/L, postmenopausal 26-110 ug/L, male 20-76 ug/L. Serum CTX 0.1-0.5ug/L, Osteocalcin 6.8-32.2ug/L. Sclerostin <80 pmol/L.

**Figure 1:**
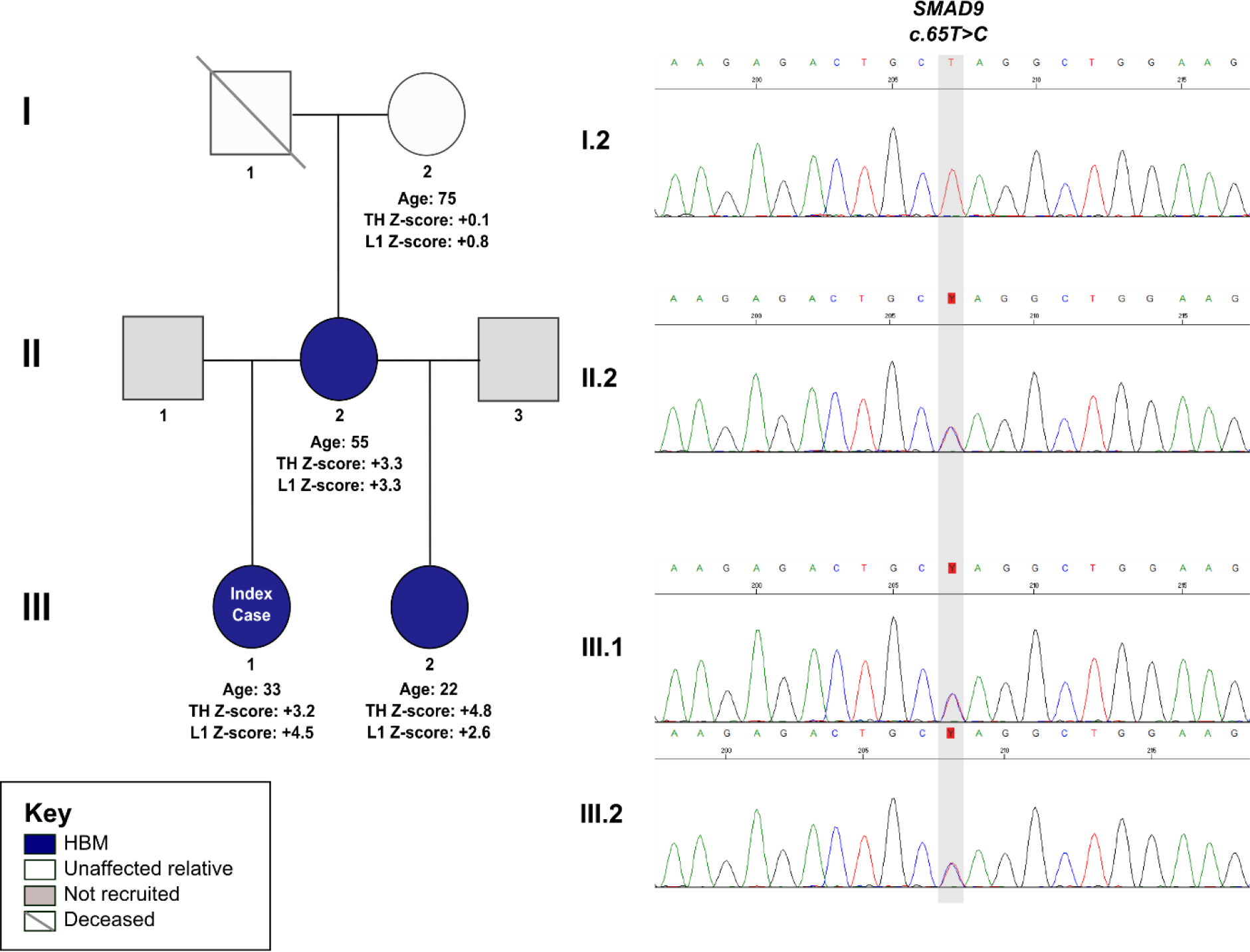
The HBM pedigree and electrophoretogram images of a segregating *SMAD9* c.65T>C, p.Leu22Pro variant.

#### Clinical phenotype (Table 1)

Clinical phenotypes of the UK family individuals are shown in Table 1; extended clinical histories are provided in Supplementary Results 1. In summary, affected individuals had high BMD Z-scores, and very high body mass index (BMI); and had not have any adult fractures, nerve compression or dental problems; however, bone pain was common, without a clear cause. There was no clinical history of intellectual impairment, pulmonary hypertension, vascular hypertension, haematological abnormalities, pubertal delay or other clinical conditions. None had been exposed to anabolic or antiresorptive medications.

#### III.1: Index Case (c.65T>C, p.Leu22Pro)

The 33-year-old index case, with BMD Z-Scores +3.2 at the total hip and +4.5 at L1, had only sustained one traumatic fracture aged 20 months. She reported lower leg and ankle pain. She was tall (>97^th^ centile) and obese, with increased shoe size, a broad frame, enlarged mandible and a 4mm torus mandibularis. She had normal joints. Radiographs showed increased cortical thickness and new bone formation at the anterior inferior iliac spines bilaterally (Supplementary Figure 1).

#### II.2: Mother of the index case (c.65T>C, p.Leu22Pro)

The 55-year-old affected mother, with BMD Z-Scores +3.3 at the total hip and at L1, had never sustained a fracture. Six years earlier she had had a right calcaneal spur surgically removed. She had widespread pain with a diagnosis of fibromyalgia. She was tall (97^th^ centile) and obese, with above average shoe size, a broad frame, enlarged mandible but no tori. She had a full range of movement in all joints, bilateral knee crepitus and bilateral pes planus.

#### III.2: Half-sister to index case (c.65T>C, p.Leu22Pro)

The 22-year-old affected half-sister, with BMD Z-Scores +4.8 at the total hip and +2.6 at L1, had not fractured. She had had sciatica for five years, lumbar back pain and fronto-temporal headaches for 11 years, with a diagnosis of migraine. She was tall (93^rd^ centile) and obese, with above average shoe size, a broad frame, enlarged mandible, a torus palitinus in the midline of her hard palate (3cm x 7mm) and normal joint movement.

#### I.2: Grandmother of index case (wild-type)

The 75-years-old grandmother, who did not have HBM, had also never sustained a fracture. She had widespread osteoarthritis and on examination had reduced extension of the right elbow and left knee, and bilateral knee crepitus. However, in contrast to other family members, she was overweight with normal shoe size, a normal frame, mandible and no tori.

#### Sequencing of pedigree

Whole exome sequencing (WES) identified a heterozygous missense variant in *SMAD9 (SMAD family Member 9 referring to homologies to the Caenorhabditis elegans SMA (small worm phenotype) and Drosophila MAD (“Mothers Against Decapentaplegic“))* (NM 001127217: exon2: c.65T>C, p.Leu22Pro), segregating with HBM (*i.e*. present in all three individuals with HBM (*III.1, II.2, III.2*), but absent from *I.2* (Figure 1). This variant (rs111748421) is rare (Exome Aggregation Consortium [ExAC] minor allele frequency [MAF] 0.0014), affects a highly evolutionarily conserved base (genomic evolutionary rate profiling [GERP] 5.53) and is predicted to be pathogenic by multiple protein-prediction algorithms (deleterious by SIFT ^12^, probably damaging by Polyphen ^13^, and disease causing by MutationTaster ^14^ and PMut ^15^).

A novel variant in *CHRNA1 (cholinergic receptor, nicotinic, alpha 1)* (c.560T>C, p.Leu187Pro) was also identified (GERP 5.29). Mutations in *CHRNA1* have been associated with congenital myasthenic syndromes (OMIM#100690), not present in this pedigree.

No variants were identified when applying a compound heterozygous or an autosomal recessive inheritance model.

### Sequencing of other HBM cases identifies two further isolated HBM cases harbouring a p.Leu22Pro mutation

WES of a further 366 HBM cases (240 isolated cases from the UK cohort with a total hip (TH) or first lumbar vertebra (L1) Z-score ≥+3.2 and 126 individuals from the Anglo-Australasian Osteoporosis Genetics Consortium (AOGC) ^16^ with either a total hip and/or lumbar spine (LS) Z-score between +2.5 and +4.0) (Supplementary Figure 2) identified two individuals with the same *SMAD9* c.65T>C, p.Leu22Pro mutation. Haplotypic analysis confirmed these women were neither related to each other nor to the pedigree described above.

#### Clinical phenotype

##### Isolated HBM case (c.65T>C, p.Leu22Pro) from the UK

***(Table 1; Supplementary Figure 3; Supplementary Results 1)*.** This 55-year-old female, with BMD Z-Scores +5.0 at the total hip and +4.7 at L1, had never fractured and had no symptoms of nerve compression. Her adult left upper cuspid tooth had never erupted; wisdom teeth had been extracted for overcrowding. She had noticed her own mandible enlargement. She had a congenital astigmatism of her left eye with poor vision, and congenital bilateral pes planus. Height was normal (30^th^ centile). She was obese with a broad frame, mandible enlargement, but no tori. She had normal joints.

##### Isolated HBM case (c.65T>C, p.Leu22Pro) from Australia

This 57-year-old female, with BMD Z-Scores +3. at the total hip and +2.7 at L1, reported a nose fracture as a child. Height was on the 45^th^ centile and she was overweight. She did not have any history of conditions affecting bone health; and had not received antiresorptive or anabolic medications. No further clinical details were available.

### Tibial pQCT evaluation

All members of the HBM pedigree, plus the additional isolated HBM case from the UK underwent pQCT scanning of the tibia (Table 2, Supplementary Table 1, Figure 2). To set these findings in context, the mean (SD) values from the four p.Leu22Pro *SMAD9* HBM cases were compared against values from 76 unrelated female HBM cases (without *SMAD9, LRP5, LRP4, or SOST* mutations) and 32 female family controls with normal DXA-measured BMD who had had pQCT scans following the same protocol ^17^. In addition to greater bone size (cross-sectional bone area) at the mid tibia (66% site), the four HBM cases with *SMAD9* mutations had greater trabecular and cortical density, cortical area and thickness and predicted bone strength (strength stain index [SSI]) than other HBM cases, and, to a greater extent, than unaffected family controls (with normal DXA-measured BMD (Table 2)). Muscle size (cross-sectional area) was also notably larger in in the *SMAD9* HBM group (Figure 2).

**Figure 2.**
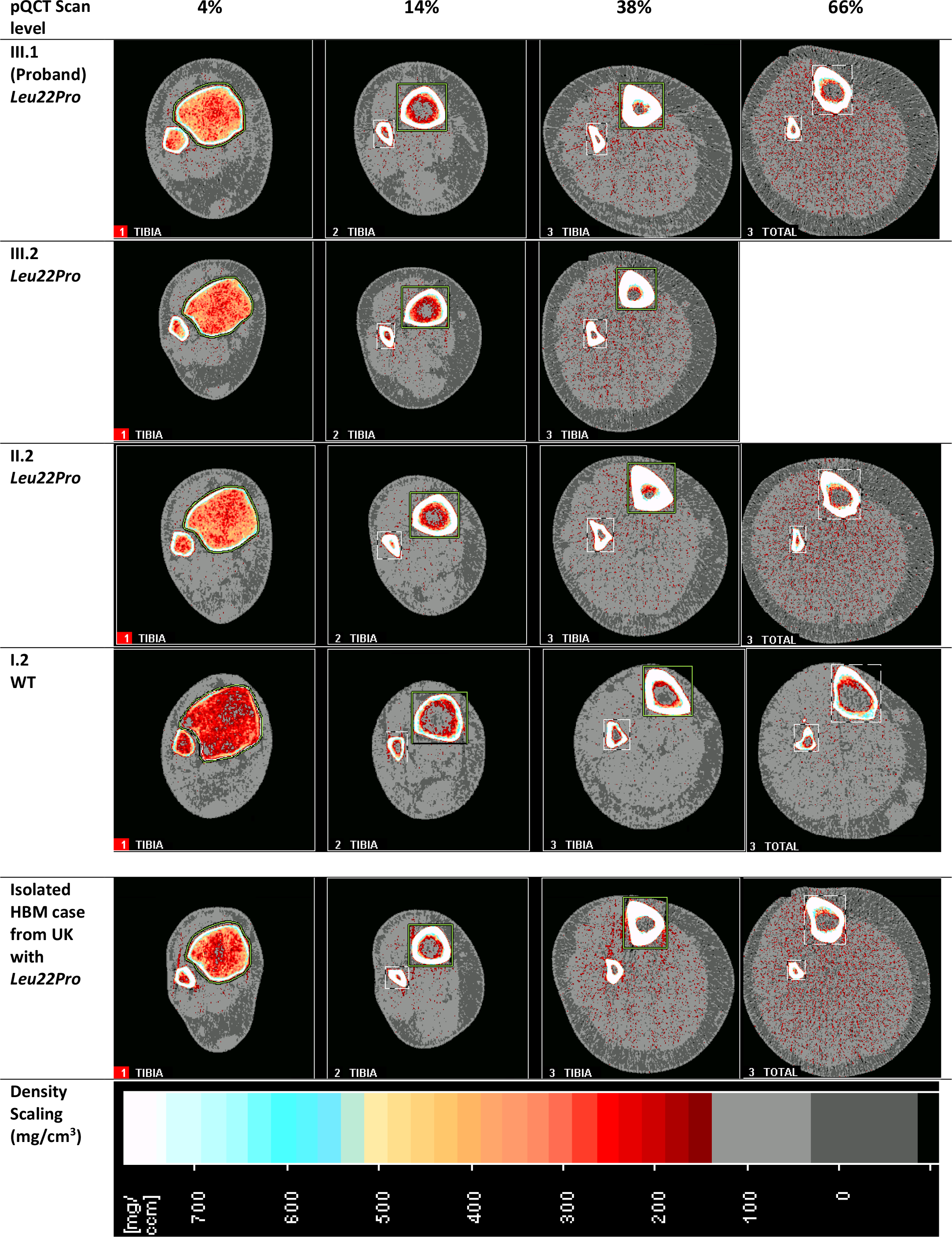
Tibia pQCT scan results from HBM pedigree and isolated HBM case with *Leu22Pro* mutation. Images from sequential pQCT scans taken at 4, 14, 38 and 66% along the tibia from the distal endplate. The 4% site shows predominantly trabecular bone, whilst cortical bone is thickest at the 38% site. The 66% site shows muscle size; of note *III.2* had too large a calf to fit within the pQCT gantry and hence this image was not attained.

**Table 2:**
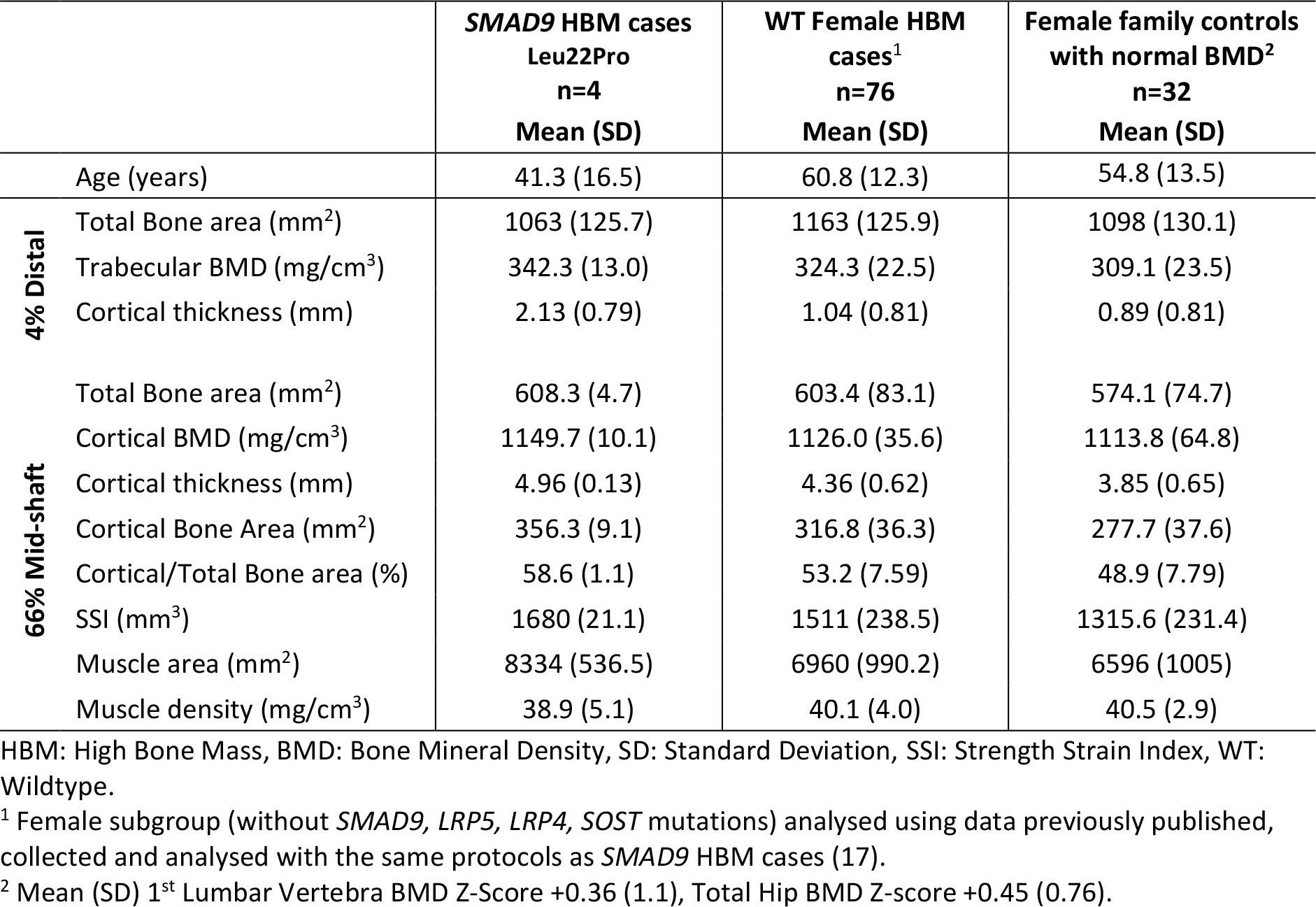
Distal and mid-shaft tibial pQCT measures in High Bone Mass cases compared with female HBM cases without *SMAD9, LRP5, LRP4, SOST* mutations, and female family controls with normal BMD.

### Sequencing of Low Bone Mass (LBM) cases

WES data from 473 women with LBM from the AOGC consortium with TH Z-scores between −1.5 and −4.0 and a LS Z-score ≤-0.5, obtained using similar methodology to the AOGC HBM cases, was interrogated (Supplementary Figure 2). The p.Leu22Pro *SMAD9* mutation was not observed.

### Common *SMAD9*-associated genetic variants and BMD

Publicly available data from a recent population-based genome-wide association study (GWAS) of eBMD (estimated BMD by heel ultrasound in the UK-Biobank study) ^18^ were used to investigate variants surrounding both *SMAD9* and *CHRNA1*. Regional association plots suggested that SNPs intersecting *SMAD9* are strongly associated with eBMD (lead SNP rs12427846 [MAF 0.25], β 0.02, SE 0.002, p=5.5×10^−16^; Figure 3). In contrast, SNPs surrounding *CHRNA1* were not robustly associated with eBMD (Supplementary Figure 4). These observations were further supported by gene-based tests of association performed in-house using 362,924 unrelated white British subjects from the UK-Biobank Study. Specifically, *SMAD9* was more strongly associated with eBMD (PJOINT = 7.94 × 10^−17^), when compared to neighbouring genes within +/− 800kb (P > 2.4 × 10^−2^) (Supplementary Table 2). No such enrichment was seen for *CHRNA1* (Supplementary Table 3). Further investigation of rs12427846 in the UK-Biobank study identified weak associations with body weight (β −0.14, SE 0.04, p=1.6×10^−3^) and with height (β −0.07, SE 0.03, p=3.5×10^−3^) with effects in the opposite direction from that seen with eBMD; however, adjustment for weight and height did not attenuate the strong association between rs12427846 and eBMD reported above. Using PhenoScanner version 2 ^19^, a PheWAS for the rs12427846 variant further identified a weak association with forearm BMD with consistent direction of effect (Supplementary Table 4). Interrogating the GWAS Catalog (https://www.ebi.ac.uk/gwas/), no associations with any trait (neither bone-related nor any other) have been reported previously for the rare (MAF=0.0014) variant rs111748421.

**Figure 3.**
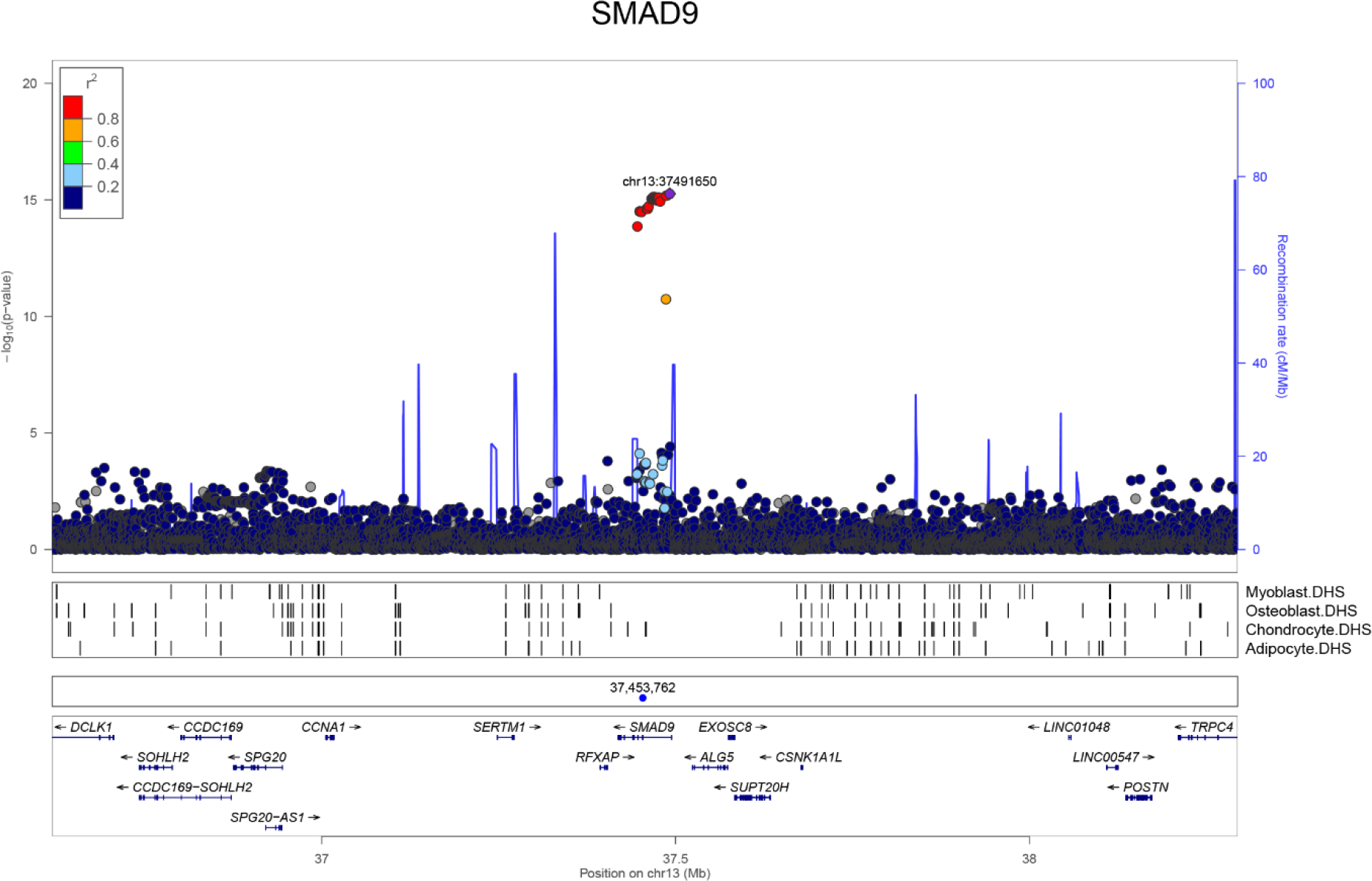
GWAS for eBMD measured in UK Biobank: Regional association plot for the locus containing *SMAD9*. Top panel: circles show unconditioned GWAS *P*-values and genomic locations of imputed SNPs within +/− 800kb of the 5’ and 3’ UTR of each gene. Different colours indicate varying degrees of pairwise linkage disequilibrium between the lead eBMD-associated SNP (rs12427846, purple diamond) and all other SNPs. Second panel: Vertical shaded areas correspond to locations of DNase I hypersensitive sites (DHSs) characteristic of: skeletal muscle myoblasts cell line (E120), osteoblast primary cells (E129), mesenchymal stem cell derived chondrocyte cultured cells (E049) and mesenchymal stem cell derived adipocyte cultured cells (E023). Red shading depicts intersections between DHSs and genome-wide significant SNPs. Black shading denotes instances in which any other SNPs intersect DHSs. Third panel: Blue circle shows the position of the putative causal variant c.65T>C, p.Leu22Pro. Fifth panel: Horizontal lines represent genes with vertical lines annotating the location of exons. Arrows indicate the direction in which each gene is transcribed.

### Smad9 expression in murine osteocytes

We next determined whether *Smad9* and *Chrna1* are expressed in osteocytes, the master cell regulators in the skeleton and key regulators of bone mass ^20^, and enriched in osteocytes compared to other cells in bone ^21^. Smad9 mRNA was highly expressed in murine osteocytes whilst Chrna1 was not (Supplementary Table 5).

### Smad9 expression in zebrafish skeletal tissue

We also examined Smad9 protein expression in the developing zebrafish skeleton ^22^ at 6- and 7-days post fertilisation (dpf) (Figure 4A). A focus of Smad9 expression was observed at the dorsal tip of the opercle, an intramembranous bone overlying the gills, adjacent to but distinct from a region of bone morphogenetic protein (BMP) reporter activity (Figure 4B). The opercula muscle group also showed evidence of BMP reporter activity, whereas Smad9 expression at this site was absent. Smad9-expressing cells in the opercle were negative for the osteoblast marker, sp7 (osterix), suggesting they are likely to represent pre-osteoblasts (Figure 4C and Supplementary video 1). Equivalent findings were observed in the branchiostegal ray bones and in the notochord at 6- and 7- dpf (Supplementary Figure 5).

**Figure 4.**
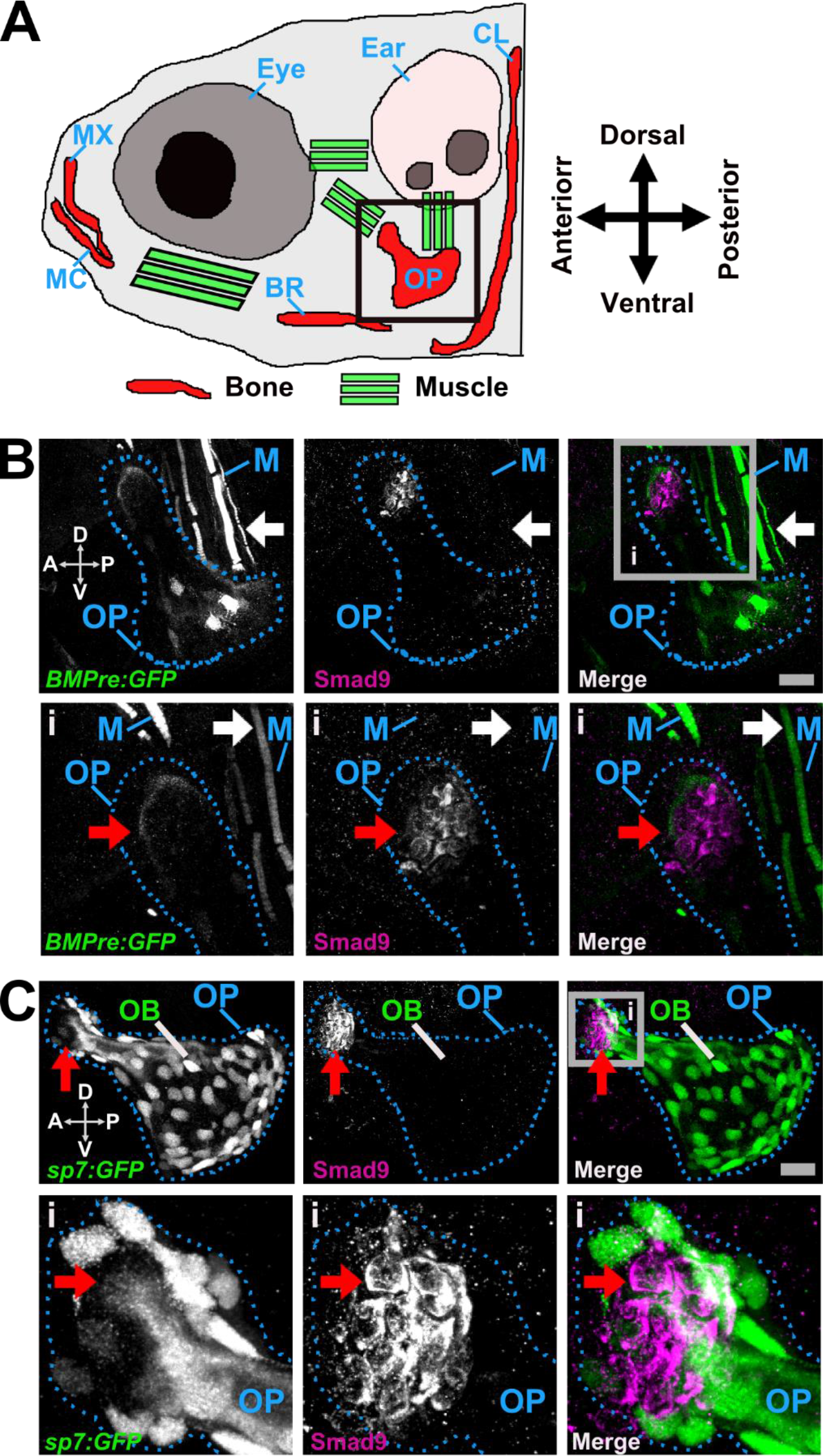
Smad9 protein expression in the larval zebrafish opercle bone. ***A. Schematic representation of the larval zebrafish head*** (6 days post fertilisation (dpf), lateral view), showing visible ossified elements (red) and the main muscle groups (green) that are green fluorescent protein (GFP) positive under control of the BMP responsive elements promoter (BMPre) transgenic reporter line (*BMPre:GFP*). The black box indicates the location of the intramembranous opercle bone as shown in panels B and C. ***B. Distinct tissue distribution of Smad9- and BMP-expressing cells*** (7dpf). Upper left panel: *BMPre:GFP* positive cells (white) in the levator operculi muscle group (white arrow) and ventral (V) side of the opercle (OP; dotted blue outline); upper middle: distinct group of Smad9 positive cells (white) in the dorsal (D) tip of the opercle; upper right: merged view showing distinct tissue expression of *BMPre:GFP* positive cells (green) and Smad9 positive cells (purple); lower left: grey box inset (i) showing faint cap of *BMPre:GFP* positive cells at the dorsal tip of the opercle (red arrow); lower middle: cluster of Smad9 positive cells; lower right: merged view confirming non-overlapping distribution of *BMPre:GFP* positive cells and Smad9 positive cells. ***C. Distinct tissue distribution of Smad9- and osterix (Sp7)-expressing cells*** (6dpf). Upper left panel: *Sp7:GFP*-positive osteoblasts (OB; white) within the opercle; upper middle: Smad9 positive cells (white) in the antero (A)-dorsal tip of the opercle (red arrow); upper right: merged view showing separation of *Sp7:GFP* positive cells (green) and Smad9 positive cells (purple) (Supplementary Video 1); lower left: the inset (i, grey box) shows few *Sp7:GFP*-positive osteoblasts within the dorsal tip of the opercle; lower middle: cluster of Smad9 positive cells; lower right: merged view confirming non-overlapping distribution of *Sp7:GFP* and Smad9 positive cells. B and C: scale bar is 20 µm, and all are maximum intensity z-projection confocal images. Abbreviations: A, anterior; BR, branchiostegal ray; CL, cleithrum; D, dorsal; M, muscle; MC, Meckel’s cartilage; MX, maxilla; OB, osteoblast; OP, operculum; P, posterior; V, ventral.

### SMAD9 protein structural modelling

SMAD9 is a TGF-beta family member DNA binding transcription factor. Phosphorylation by BMP-ligand-bound type 1 receptor kinase activates SMAD9, which translocates from the cytoplasm to the nucleus to regulate target gene expression ^23^. The seven exons of human *SMAD9* encode a protein of 467 amino acids that contains two MAD-homology (MH) domains (MAD: Mother against Dpp) separated by a linker region (Figure 5). The p.Leu22Pro *SMAD9* mutation is located within the MH1 domain responsible for DNA-binding (Figure 5), and lies in the hydrophobic face of the N-terminal alpha helix (helix-1) (Figure 6, Supplementary Video 2). Helix-1 packs against a groove made by helix-2 and −3 within MH1, forming part of the hydrophobic core of this domain. Substitution of leucine by proline will: a) introduce a less hydrophobic residue into this position; and b) compromise the α-helical fold by disrupting the canonical hydrogen bonding of helix-1. Thus, modelling suggests that this mutation will disrupt the MH1 domain so severely that SMAD9 can no longer bind DNA and/or will be unstable leading to protein degradation.

**Figure 5.**
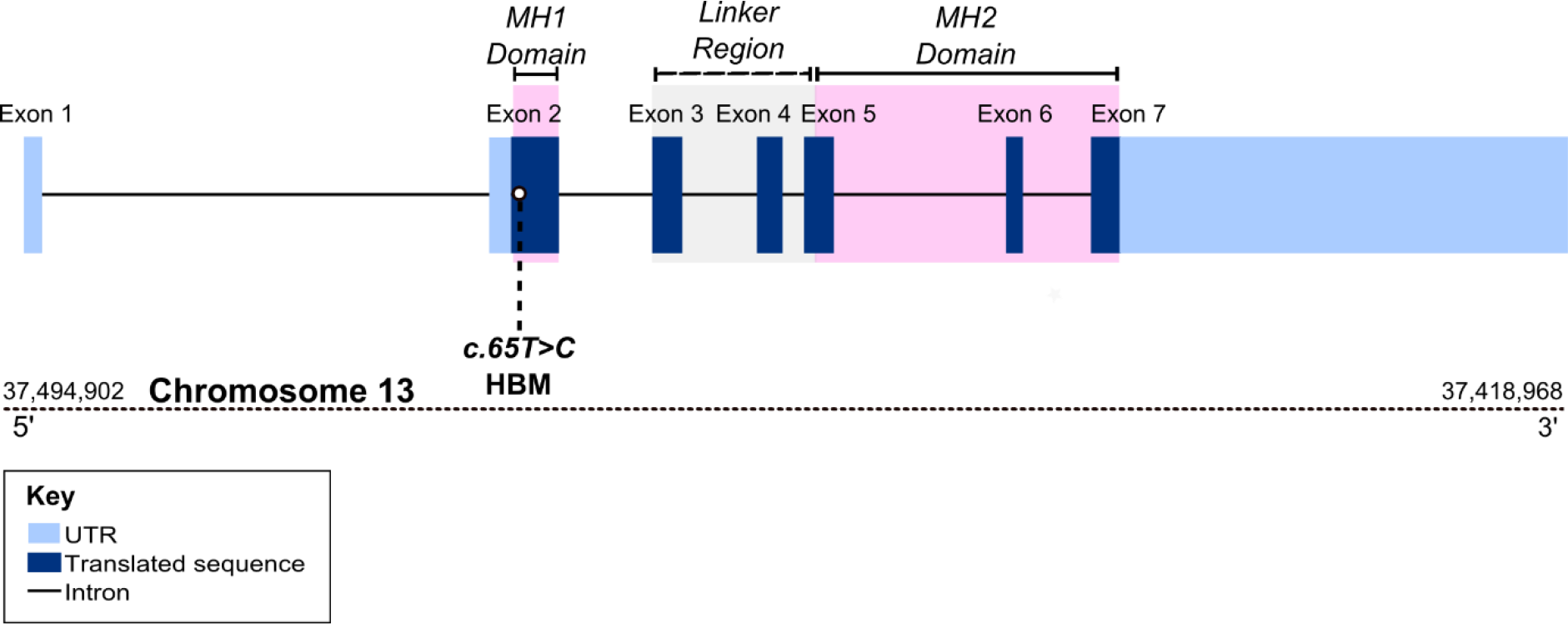
Figure illustrating position of c.65T>C p.Leu22Pro variant within *SMAD9*.

**Figure 6.**
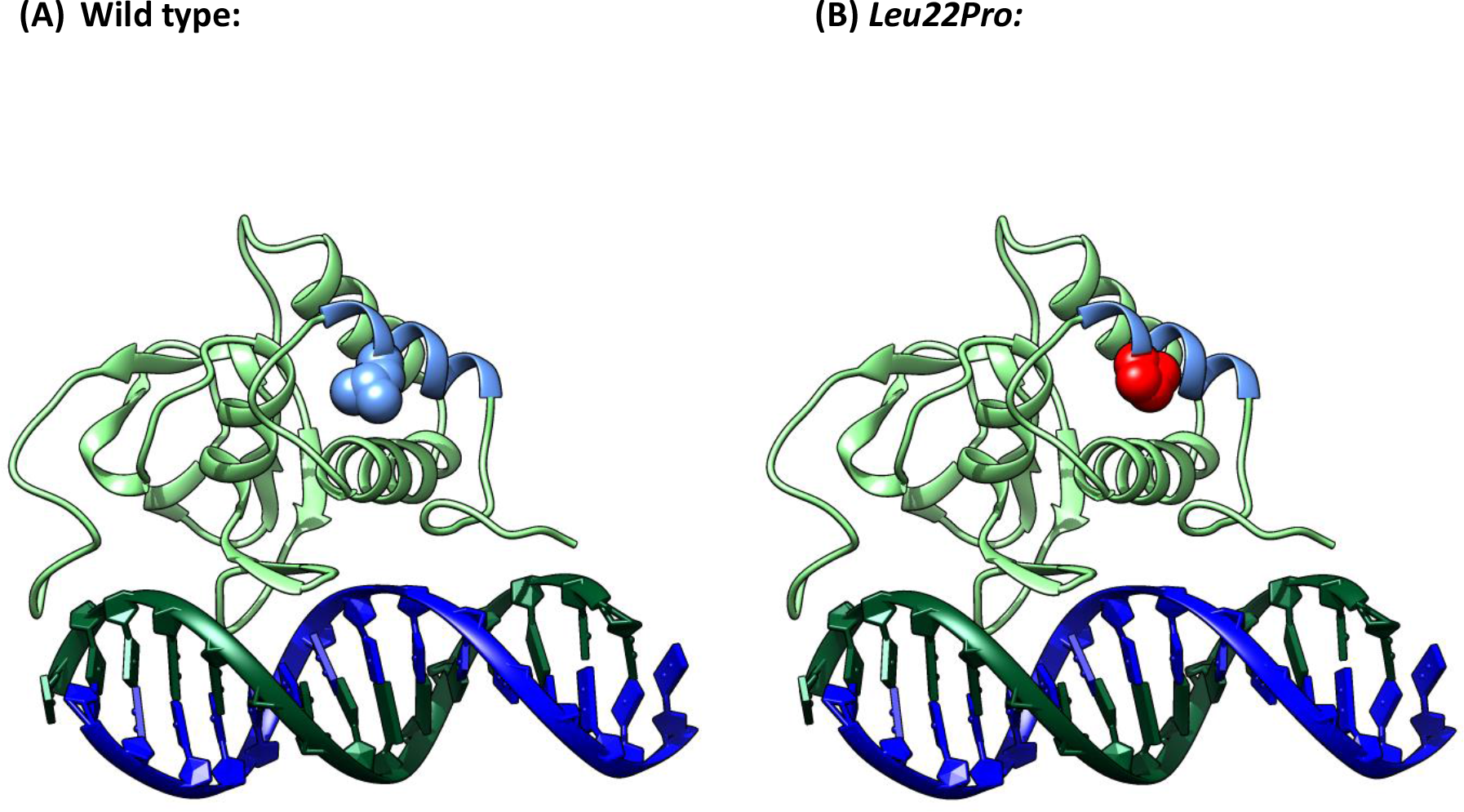
(A) Wildtype (WT) *SMAD9* MH1 domain (light green ribbon with helix-1 in light blue) binding the DNA helix (dark blue/dark green). L22 is shown in blue space-filling. (B) L22P, shown in red space-filling, is predicted to severely disrupt the structure of the MH1 domain. Supplementary Video 2: 3-dimensional rotating image

## Discussion

We report the first HBM pedigree with a segregating *SMAD9* mutation, with replication in two further unrelated individuals with HBM. *SMAD9* (also known *SMAD8, MADH6,* and *MADH9*) encodes a downstream modulator of the BMP signalling pathway. BMPs, members of the TGF-β superfamily, were first reported in 1977 to induce the formation of bone and cartilage when transplanted subcutaneously ^24^. SMADs, activated by ligand-binding of cell surface BMP receptors, mediate down-stream intracellular signalling and biological responses induced by BMPs ^25^. Smad6 and Smad7 both inhibit BMP receptor activation and downstream signalling, as does Smad9 by more direct transcriptional repression ^23^. Our *in-silico* protein modelling predicts that the p.Leu22Pro mutation severely disrupts the structure of the MH1 DNA binding domain of *SMAD9*, leading to loss of function.

Whilst few previous studies have examined sites of Smad9 tissue expression, we have confirmed that Smad9 is expressed in both mouse cortical bone derived osteocytes, and in skeletal elements of zebrafish larvae. Moreover, we observed that BMP reporter activity in zebrafish was absent at sites of Smad9 expression, consistent with a functional role in BMP repression ^23^. Taken together our findings suggest that *SMAD9* c.65T>C is a loss-of-function mutation, causing HBM through a novel mechanism of enhanced bone formation as a consequence of reduced BMP inhibition.

Further, we have shown that the region containing *SMAD9* is strongly associated with BMD within the general population. Common variants intersecting *SMAD9* associate with population-based measures of eBMD, as evidenced recently ^18,26^ and from our gene-based tests of association presented here. Furthermore, rs12427846 [the lead SNP from these eBMD results] is associated with DXA-measured total body BMD ^27^ and fracture risk ^18^. These findings, which provide further evidence of the importance of *SMAD9* in bone biology, are equivalent to reported associations for common variants annotated to *LRP5* and *SOST* genes, both similarly implicated in monogenic HBM disorders ^28,29^.

We have previously estimated unexplained HBM to have a prevalence of 0.181% amongst a DXA-scanned adult population in the UK ^10^. As two of 248 cases fulfilling our stringent HBM phenotype definition (Supplementary Figure 2) were found to harbour the p.Leu22Pro mutation, we would estimate *SMAD9* HBM to have a prevalence of approximately 1 in 100,000 (1.46 x 10^−5^); less common than *LRP5* HBM ^9^.

The clinical phenotype of p.Leu22Pro *SMAD9* HBM includes mandible enlargement, a broad frame, torus palitinus, pes planus, increased shoe size and a tendency to sink when swimming. Adult fractures were not reported, raising the possibility of increased skeletal strength which would be supported by greater cortical bone and an increased strength-strain index (SSI) quantified by pQCT (discussed further below), both of which promote fracture resistance. Mandible enlargement, torus palitinus, a tendency to sink when swimming, and an absence of adult fractures are reminiscent of *LRP5* HBM ^7,30^. Encouragingly, unlike sclerosteosis (due to anabolic *SOST* mutations) and some *LRP5* HBM cases ^9,31^, nerve compression does not seem to be a feature of *SMAD9* HBM.

The musculoskeletal phenotype of p.Leu22Pro *SMAD9* HBM includes high BMD Z-Scores (+3 to +5), with increased fat and lean mass. pQCT revealed that both volumetric cortical and trabecular bone density are increased. Furthermore, the larger bone size with greater cortical thickness and cortical area, suggest a possible combination of enhanced modelling to increase periosteal expansion and reduced bone remodelling to reduce endosteal expansion. In support, bone turnover markers are at the lower end of the normal range. This phenotype also mimics that previously described for human *LRP5* HBM ^32^. Plasma sclerostin is not elevated, in contrast to *LRP5* HBM ^33^, suggesting that a negative feedback loop downregulating WNT signalling is not present.

pQCT further confirmed that larger muscle size is another phenotypic feature of *SMAD9* HBM, with an increase in muscle cross-sectional area. This contrasts with the suggestion from our zebrafish studies that Smad9 expression in muscle tissue may be limited. Given the well-recognised cross-talk between muscle and bone ^34^ and the large BMI of these individuals, it is conceivable that the increase in muscle size is a consequence of increased bone mass. However, similar increases in muscle size have not been reported in other monogenic HBM conditions (*i.e. LRP5* or *SOST* HBM) with equivalent increases in BMD.

Heterozygous truncating *SMAD9* mutations are associated with primary pulmonary hypertension (OMIM#615342) ^35^, a phenotype not apparent in our HBM cases. Reported mutations affect a different domain from the mutation observed here (p.Leu22Pro), with p.Cys202X ^35^ and p.Arg294X ^36^ truncating the SMAD9 protein in the linker region between MH1 and MH2. A truncating mutation (p.Arg247X) has been associated with cerebral arteriovenous malformations in childhood ^37^. An activating heterozygous p.Val90Met germline mutation, affecting the 4^th^ α-helix of MH1 and close to the DNA binding interface, has been described in one pedigree with hamartomatous polyposis ^38^. In contrast to p.Leu22Pro, p.Val90Met appears to be a gain-of-function mutation, thought to arise from a steric clash, prompting a His104 residue to enhance DNA binding ^38^. Such examples of diverse phenotypes arising from mutations in differing exons of the same gene are well recognised, *e.g*. differing mutations in *FBN1 (Fibrillin 1)* can cause Marfan syndrome (with associated tall stature) (OMIM#154700), acromicric dysplasias (with short stature) (OMIM#102370), or stiff skin syndrome (OMIM#184900) ^39-41^. Similarly mutations in *PIGN (Phosphatidylinositol Glycan Anchor Biosynthesis Class N)* are associated with multiple congenital anomalies-hypotonia-seizures syndrome 1 (OMIM#614080) ^42^ and Fryns syndrome (congenital diaphragmatic hernia and other dysmorphic features) (OMIM#229850) ^43^.

We are only aware of one other skeletal dysplasia reported in association with an inhibitory SMAD (which include *SMAD6* and *SMAD7*). A rare *SMAD6* mutation has been associated with susceptibility to non-syndromic midline craniosynostosis 7 (OMIM#617439) - but only in the context of co-inheritance of a common variant in *BMP2* strongly associated with this condition, a rare example of two locus inheritance^44^. Interestingly, amongst the 1103 conditionally independent SNPs reaching genome-wide significance in the UK-Biobank eBMD GWAS (population n=426,824), as well as identifying the *SMAD9* locus, four novel SNPs annotating to *SMAD7* were also reported (in addition to three established SNPs associated with *SMAD3*), all suggesting variation in inhibitory SMADs is likely of functional importance in human biology ^18^.

The phenotype we describe here contrasts with that of activating mutations of the BMP receptor, *ACVR1*, which increase BMP signalling. However, in contrast to p.Leu22Pro *SMAD9* HBM*, ACVR1* mutations lead to a fatal condition, Fibrous Ossificans Progressiva (FOP, OMIM#135100) ^45^. In FOP, muscle tissue differentiates into bone following trivial injury, resulting in the formation of mature bone at multiple extra-skeletal sites. *ACVR1* mutations may produce a more severe phenotype, compared with loss-of-function mutations in *SMAD9* reported here, since *ACVR1* also activates non-SMAD dependent BMP signalling cascades such as the NF-κB and p38 MAP kinase (p38MAPK) pathways, which are upregulated in FOP *ACVR1* R206H monocytes ^46^.

In view of the benign phenotype which we observed in p.Leu22Pro *SMAD9* carriers, our findings suggest that SMAD9 is worthy of consideration as a drug target for osteoporosis. Our zebrafish studies suggest that Smad9 is expressed in pre-osteoblasts, consistent with the profile of an anabolic target capable of stimulating new bone formation through recruitment of early osteoblast progenitors. Given the pathological consequence of excess BMP activation in FOP, this pathway has not previously been prioritised as a possible target in osteoporosis research, despite the profound bone anabolic potential. Interestingly, phosphorylation of Smad9, as part of the Smad1/5/9 heterotrimer, has been researched in relation to fracture healing and bone regeneration: G-protein-coupled receptor kinase 2-interacting protein-1 (GIT1), a shuttle protein in osteoblasts, has been shown to regulate Smad1/5/9 phosphorylation which then mediates BMP2 regulation of Runx2 expression and thus endochondral bone formation at the site of fracture ^47,48^. Moreover, local BMP administration has been used to promote bone repair following surgery ^49^. Based on our findings, it is tempting to speculate that treatments intended to suppress SMAD9 activity might prove useful in treating osteoporosis, fractures, and possibly also sarcopenia.

Our study has limitations. All individuals with p.Leu22Pro *SMAD9* HBM were female, which reflects the study design which favoured those with a historical DXA scan who are more likely to be female. Whether findings will be similar in men is unknown, although no sex-gene interaction has been described for the *LRP4, LRP5, LRP6,* or *SOST* sclerosing bone dysplasias. In the recent UK-Biobank eBMD GWAS, LD score regression analyses suggested that the genetic architecture influencing male and female eBMD was largely shared but that there were some significant differences between the sexes (rG=0.91, SE=0.012, p<0.001) ^18^, consistent with earlier epidemiological studies ^50^. The small sample of *SMAD9* HBM cases (n=4 with pQCT) limited our ability to robustly evaluate associations statistically. The p.Leu22Pro mutation is a reported SNP with an rsID, indicating this is carried within the general population (*e.g*. in the UK at estimated 92,428 people might be expected to carry this mutation). This may be the case, given there is no indication that the phenotype affects reproductive fitness, and none of the phenotypic characteristics are deleterious to health; and HBM will not be overt unless a DXA scan is performed. The absence of reported association for rs111748421 in the GWAS catalogue is not surprising given the difficulty of imputing extremely rare variants/ mutations for analysis in GWAS. Our GWAS was based on estimated heel BMD quantified by ultrasound rather than DXA-measured BMD. Estimated heel BMD is not used routinely in clinical practice. However, we have previously demonstrated a strong overlap between genetic loci identified by eBMD GWAS and by DXA-measured BMD GWAS ^26^.

## Conclusions

We report *SMAD9* as a novel HBM-causing gene. The clinical phenotype of p.Leu22Pro *SMAD9* HBM has many features in common with that of *LRP5* HBM, but lacks the deleterious features which characterise *SOST* HBM (sclerosteosis). As has been reported for both *LRP5* and *SOST*, we demonstrate that *SMAD9* can be disrupted by a single point mutation causing an extreme bone phenotype and perturbed by common variation affecting bone density within the general population. The role of SMAD9 in bone biology is supported by our finding of high levels of Smad9 expression in murine osteocytes, and in skeletal elements of zebrafish larvae. Smad9 is thought to inhibit BMP signalling to reduce osteoblast activity; thus, we hypothesise *SMAD9* c.65T>C is a loss-of-function mutation reducing BMP inhibition, ultimately leading to enhanced bone formation. Our findings support SMAD9, and its role within the SMAD9-dependent BMP signalling pathway, as a potential novel anabolic target for osteoporosis therapeutics which warrants further investigation.

## Methods

### The UK HBM Cohort

The HBM study is a UK-based multi-centred observational study of adults with unexplained HBM, identified incidentally on routine clinical DXA scanning. Full details of DXA database screening and participant recruitment have previously been reported ^10^. In brief, DXA databases containing 335,115 DXA scans were initially searched for a BMD T or Z-score ≥+4 at any site within the LS or hip, at 13 UK centers. All 1505 DXA images were visually inspected; 962 cases with established and/or artefactual causes of raised BMD were excluded (including exclusion of artefact from LS osteoarthritis (see Supplementary Methods 1). As a generalized HBM trait should affect both spine and hip BMD, though not necessarily equally, HBM was defined as a) L1 Z-score of ≥ +3.2 plus TH Z-score of ≥ +1.2 and/or b) TH Z-score ≥ +3.2 plus L1 Z-score of ≥ +1.2 (using age and gender-adjusted BMD Z-scores). A threshold of +3.2 was in keeping with the only published precedent for identifying HBM using DXA ^7^. Z rather than T-score was used to limit age bias.

#### Recruitment

533 unexplained HBM index cases were invited to participate. 248 (47%) were recruited ^10^. Index cases were asked to pass on study invitations to their first-degree relatives and spouse/partner(s). These relatives and spouses were invited only once, and non-responders were not followed-up.

Relatives/spouses with HBM were in turn asked to pass on study invitations to their (previously uninvited) first-degree relatives and spouses. 893 relatives were invited to participate, of whom 236 (26.4%) were recruited. 217 spouses/ partners were invited to participate, of whom 61 (28.1%) were recruited; 2 individuals invited 2 partners ^10^. All participants were clinically assessed using a standardised structured history and examination data collection tool.

#### DXA measurements

Participants underwent DXA scanning (repeat in the case of index cases) using either GE Lunar Prodigy DXA (software version 13.2, GEHealthcare, Madison, WI, USA) or Hologic Discovery/W DXA (software version Apex 3.0, Hologic Inc. Bedford, MA, USA). Scans were acquired and analysed according to each manufacturer’s standard scanning and positioning protocols as previously described ^51^. Total Body (TB) bone mineral content (BMC) and density (BMD), Fat Mass (FM) and Lean Mass (LM) were measured, together with L1 and total hip BMD. All DXA images were reviewed for quality control purposes.

#### Peripheral Quantitative Computed Tomography (pQCT)

pQCT scans were performed at the distal and mid-shaft of the tibia (4, 14, 38 & 66% from the distal endplate) in the non-dominant lower limb using a Stratec XCT2000L (Stratec Medizintechnik, Pforzheim, Germany); voxel size 0.5 mm, CT speed 30 mm/second, XCT software version 5.50d; as published previously ^17^. In brief, a reference line at the distal endplate was determined from initial frontal scout view. Cortical bone was defined using a threshold above 650mg/cm^3^ (optimal for bone geometry ^52^). Trabecular bone was identified by elimination of cortical bone and therefore trabecular bone mineral density (tBMD) was defined as a density <650mg/cm^3^. Cortical thickness, periosteal circumference and endosteal circumference were derived using a circular ring model.

Further cortical parameters were measured: cortical bone mineral density (cBMD), total bone area (TBA) (*i.e*. total bone cross-section, reflecting periosteal expansion), cortical bone area (CBA) (reflecting a combination of periosteal and endosteal expansion) and CBA/TBA (%). Strength Strain Index (SSI) was calculated according to Stratec’s user manual (SSI= SM*(cBMD[mg/cm^3^]/ 1200[mg/cm^3^]), where 1200 mg/cm^3^ represents the normal physiological density of bone (stated by Stratec) and SM (Section Modulus)=CSMI/ periosteal radius, where CSMI (Cross-Sectional Moment of Inertia [cm^4^])=Π(periosteal radius^4^-endosteal radius^4^)/4) ^53^. Twenty population controls were scanned twice on the same day after repositioning; measurement precision (CV) was typically 1 to 3 % [11]. Stratec pQCT machines were calibrated using a COMAC phantom; mean (SD) difference between scanners was 1.18 (0.82) %.

#### Blood testing including bone turnover markers and DNA extraction

Two non-fasted EDTA samples were collected and serum separated and frozen within 4 hours to − 80°C. Bone formation (Procollagen type 1 amino-terminal propeptide [P1NP], total osteocalcin) and resorption (β-C-telopeptides of type I collagen [βCTX]) markers were measured by Electrochemiluminescence immunoassays (ECLIA) performed on the COBAS e601 analyser (Roche Diagnostics, Burgess Hill, UK). Inter-assay coefficient of variation (CV) for P1NP was <3.0% across the range between 5-1200 µg/L; osteocalcin was <5.0% across the range between 0.5-300 µg/L; βCTX was <3.0% across the range between 0.01-6 µg/L. Sclerostin was measured using an enzyme-linked immunosorbent assay (ELISA) kit BI-20492 (Biomedica GmbH, Vienna, Austria) with assay CV of <9.0% across the range 2.6-240 pmol/L. DNA was extracted from peripheral venous blood using standard phenol/ chloroform extraction.

#### Exclusion of known monogenic causes of HBM

Sanger sequencing of all HBM index cases for exons 2, 3 and 4 of *LRP5*, *SOST* (including the van Buchem disease deletion) and *LRP4* (exons 25 and 26) excluded seven with *LRP5* mutations and one with a *SOST* mutation ^9^, leaving 240 unexplained HBM individuals.

#### Ethics approval

Written informed consent was collected for all participants in line with the Declaration of Helsinki ^54^. This study was approved by the Bath Multi-centre Research Ethics Committee (REC: 05/Q2001/78) and at each NHS Local REC.

### Anglo-Australasian Osteoporosis Genetics Consortium (AOGC) HBM and LBM cases

The original AOGC extreme truncate population included 1128 Australian, 74 New Zealand and 753 British women, aged between 55–85 years, five or more years postmenopausal, with either HBM (age and gender-adjusted BMD Z-scores of +1.5 to +4.0, n=1055) or LBM (age- and gender-adjusted BMD Z-scores of −4.0 to −1.5, n=900) ^55^. LBM cases were excluded if they had secondary causes of osteoporosis (as previously described ^55^). Unrelated samples of Caucasian ancestry with complete height and weight data and enough high-quality genomic DNA were available in 947 AOGC individuals (426 AOGC high and 521 AOGC low BMD), from which (due to computation capacity limiting sample size) the most extreme HBM cases were then selected, using a threshold TH or LS Z-score ≥+2.5, and the most extreme LBM cases using a LS Z-Score ≤-0.5, so 126 HBM and 493 LBM samples were chosen to undergo WES.

#### Ethics approval

The AOGC study was approved by the Queensland Office of Human Research Ethics Committee (Ref:2008/018), the University of Queensland (Ref:200800376) and/or relevant research ethics authorities at each participating centre. Directly recruited participants gave written, informed consent. Some participants were recruited through genetic and/or clinical studies (all with appropriate ethical approval) but also provided written informed consent to contribute to collaborative genetic studies ^56-58^. DNA was obtained from peripheral venous blood or from saliva through standard methods.

### Whole exome sequencing

Sequencing libraries for 859 samples (240 UK HBM, 126 AOGC HBM, 493 AOGC LBM) were constructed in two batches, determined by sample availability, the first (85 HBM, 619 AOGC) using the Illumina TruSeqDNA sample preparation kit, combined in pools of six for target capture by the Illumina TruSeq Exome v2.0 Enrichment Kit (64Mb capture) and assessed pre and post-capture for quality and yield with the Agilent High Sensitivity DNA assay and KAPA Library Quantification Kit. For the second batch (155 HBM), exome capture was performed using the Nextera^®^ Rapid Capture Exome Enrichment Kit (Cat No FC-140-1003) (62Mb capture) (Illumina, San Diego, California, USA) as per the Nextera^®^ Rapid Capture Enrichment. In both cases massive parallel sequencing was performed with six samples per flow cell lane on the Illumina HiSeq2000 platform and version 2 SBS reagents to generate 100 bp paired-end reads. Base calling, sequence alignment and variant calling were all performed as previously described ^59^, in brief, after demultiplexing, the Illumina Data Analysis Pipeline software (CASAVA v.1.8.2) was used for initial base calling. Sequence data were aligned to the current build of the human genome (UCSC Genome Browser, hg19, released February 2009) via the Novoalign alignment tool (v.2.08.02 1) ^60^; sequence alignment files were converted by SAMtools (v.0.1.14) ^61^ and Picard tools (v.1.42). SNPs and indels were called with the Genome Analysis Toolkit (GATK v.5506) ^62,63^ and annotated by ANNOVAR ^64^. Further analysis of sequence data was performed with custom scripts employing R and Bioconductor. Good-quality SNPs, excluding all those with a genotype quality score, determined by the GATK algorithm, <60 (range 0 [poor] to 100 [excellent]), were retained. Platform related artefacts variants were identified as variants where the allele count was 6 SDs higher than that expected from the maximum population MAF under the binomial approximation. Remaining SNPs and indels were assessed according to prediction of potentially damaging consequence (“nonsynonymous SNV,” “splicing,” “frameshift substitution,” “stopgain SNV,” “stoploss SNV”) by using both RefSeq and UCSC transcripts. Further filtering excluded SNPs with a minor allele frequency (MAF) <0.05 (observed in NCBI dbSNP (GRCh37/ Hg19), 1000 Genomes ^65^, ExAC (http://exac.broadinstitute.org/), or internal databases from >3000 exomes), 1000 Genomes small indels (called with the DINDEL program ^66^). Variants not present in any of these databases were considered novel. Platform related artefacts were identified as variants where the allele count was 6 standard deviation higher than that expected from the maximum population MAF under the binomial approximation. Genotype Quality scores for the two additional isolated HBM cases were 99 and read depths were 15,14 and 21,19.

### Filtering pipeline applied to unexplained HBM pedigrees

After quality-filtering as described above, data were analyzed for carriage of at least one rare (either novel or maximum population based MAF <0.005) nonsynonymous SNV or indel in a highly conserved region (GERP score <1.5) of a gene, carried by the affected individuals and not carried by unaffected individuals (*i.e*. autosomal dominant carriage model). Data were then filtered based on functional prediction of SNVs using Polyphen ^13^ to identify ‘probably damaging’ and SIFT ^12^ ‘deleterious’ SNVs. Compound heterozygous and homozygous inheritance were also assessed.

### Sanger sequencing validation of pedigree based HBM mutation

Polymerase chain reaction (PCR) amplification of identified exons was performed on 50ng genomic DNA in a reaction mix consisting; 10X Immolase reaction buffer, 10mM dNTPs, 50mM MgCl2, 5µM each primer, 0.5 U Immolase polymerase Taq (Bioline Reagents Ltd, London), and water to final volume of 25µl. PCR cycling conditions and primer sequences are shown in (Supplementary Methods 2). Samples were Sanger sequenced using standard techniques (BigDye v3.1 chemistry, Life Technologies Corporation, California), and capillary sequenced (3130 Genetic Analyzer, Life Technologies Corporation, California). Electropherograms were aligned and analysed using sequence analysis software Genalys (Version 2.0 ß, Masazumi Takahashi).

### Multi-marker Analysis of GenoMic Annotation (MAGMA) in UK Biobank

Gene-based tests of association were performed on 362,924 unrelated white British subjects (54% female, GCTA-GRM derived pairwise relatedness < 0.10) from the UK Biobank study that had valid quantitative ultrasound derived heel eBMD and high quality genome-wide HRC and 1000G/UK10K imputed data from the January 2018 release [*i.e*. 20,490,436 genetic variants with an information quality score > 0.3, MAF > 0.05, minor allele count > 5, genotyping hard call rate > 0.95, and weak evidence of deviation from Hardy-Weinberg equilibrium (p >1×10^−6^)]. A detailed description of the methodology used to generate this resource is published elsewhere ^18^. Gene-based tests of association were implemented in MAGMA v1.06 ^67^ using a multi-model approach that combines the association results from three separate gene analysis models: principal components regression, SNP-wise Mean χ2 model [*i.e*. test statistic derived as the sum of −log(SNP p-value)] and SNP-wise Top χ2 model [(test statistic derived as the sum of −log(SNP p-value) for top SNPs)] to produce an aggregate p-value corresponding to the strength of evidence of association between each of the 19,361 protein coding genes (+/− 20kb) and BMD, adjusting for age, sex, genotyping array, assessment centre and ancestry informative principal components 1 – 20. Summary results statistics for all genes within +/− 800kb of *SMAD9* and *CHNR1* were looked up. Regional association plots were generated using LocusZoom (v1.3) ^68^ in conjunction with summary association results from Morris *et al*. 2018 ^18^.

### Gene expression in murine osteocytes

Whole transcriptome sequencing data from the primary osteocytes of four different bone types (tibia, femur, humerus and calvaria) from mice (marrow removed, n=8 per bone) were analysed. A threshold of expression was determined based on the distribution of normalised gene expression for each sample ^69^. “Expressed” genes were above this threshold for all 8 of 8 replicates in any bone type. Osteocyte enriched expression of these genes in the skeleton was determined by comparing transcriptome-sequencing data from bone-samples with osteocytes isolated versus those samples with marrow left intact (n=5 per group) ^21^.

### Replication in high BMD populations

WES data from AOGC were analysed to identify any individual who carried the same rare (MAF <0.025) mutation as identified from analysis of the HBM pedigree. Polyphen ^13^ and SIFT ^12^, PMut ^15^ and MutationTaster ^14^ were used for *in silico* functional prediction. When the same point mutation was identified in more than one individual, haplotypes were compared between index case samples genotyped using an Infinium OmniExpress-12v1.0 GWAS chip read using an Illumina iScan (San Diego, California, USA), with genotype clustering performed using Illumina BeadStudio software.

### Protein structural modelling

The amino-acid sequence of human SMAD9 was passed to the HHPred server (https://toolkit.tuebingen.mpg.de/#/tools/hhpred) ^70^. This located the best template structures in the Protein Databank for the MH1 domain, 5X6G (mouse SMAD5; 92% identity), and the MH2 domain, 3GMJ (*Drosophila melanogaster* MAD; 75% identity). Modeller was used to build the domain models according to the HHPred alignments ^71^. Chimera was used to introduce point mutations and remodel the domain swapping in the SMAD9-MH1 model ^72^.

### Zebrafish studies

*BMPre:GFP* (*Tg(5xBMPRE-Xla.Id3:GFP)*) ^73^ and *sp7:GFP* (*Tg(Ola.sp7:NLS-GFP)*) ^74^ transgenic fish (in London AB background) were maintained in standard conditions ^75^. Experiments were approved by the University of Bristol Animal Welfare and Ethical Review Body (AWERB) and performed in accordance with a UK Home Office project license. Developmentally staged larvae (following euthanisation in MS222) were fixed in 4% paraformaldehyde (1 hour), dehydrated to 100% methanol and stored at −20°C prior to staining. Immunolabelling protocol was as previously described ^76^.

Primary antibodies: anti-Smad9 (rabbit polyclonal, Abcam, ab96698) used at a 1/100 dilution and anti-GFP (chicken polyclonal, Abcam, ab13970) used at a 1/200 dilution in blocking buffer (5% horse serum). Secondary antibodies were used (A21206 and A11041, Invitrogen) in a 1/400 dilution and samples were incubated with DAPI (Sigma-Aldrich, 1/1000 dilution) to visualise nuclei. Samples were mounted in 1% low melting point agarose and images were taken with a confocal laser scanning microscope (Leica, SP5II AOBS attached to a Leica DM I6000 inverted epifluorescence microscope) using a 40X PL APO CS (1.3 numerical aperture) lens. Images were processed and colour balanced in Fiji^77^.

**Supplementary Information** is available in the online version of this paper.

## Supporting information

All supplemental information

## Acknowledgements

Full acknowledgements are listed in the Supplementary Information.

## Funding

CLG was funded by the Wellcome Trust (080280/Z/06/Z), the EU 7th Framework Programme under grant agreement number 247642 (GEoCoDE), a British Geriatric Society travel grant, and Versus Arthritis (formerly Arthritis Research UK) (grant ref 20000). This study was supported by the NIHR CRN (portfolio number 5163). DB received travel grants from The Harold Hyam Wingate Foundation, the Disease Models and Mechanisms journal (DMMTF-180208), and an Elizabeth Blackwell Institute for Health Research (University of Bristol) discipline hopping fellowship via a Wellcome Trust Institutional Strategic Support Grant (204813/Z/16/Z). AH is funded by the Wellcome Trust (grant ref 20378/Z/16/Z). LP, AH and GDS work in a unit which receives UK Medical Research Council funding (MC_UU_12013/4). CH and DB were funded by Versus Arthritis (21211, 21937, 19476). AML is supported by an NHMRC Early Career Fellowship. JPK is funded by a University of Queensland Development Fellowship (UQFEL1718945) his contribution is supported by a National Health and Medical Research Council (Australia) project grant (GNT1158758). MAB is supported by an NHMRC Senior Principal Research Fellowship. PIC is funded by a Wellcome Trust Strategic Award (grant ref 101123). The AOGC was funded by the National Health and Medical Research Council (Australia) (Project Grants 511132 and 1032571).

## Author Contributions

Conception CLG, GDS, MAB, JHT, ELD Design CLG, GDS, MAB, PL, JHT, ELD

Data acquisition CLG, DB, LW, PC, SY, WF, JCYT, CH, MAB, ELD

Analysis CLG, DB, RBS, LW, AH, SY, PC, AML, CH, JPK, PL, ELD

Interpretation CLG, DB, RBS, AH, SY, PC, AML, MAB, CH, JPK, PL, JHT, ELD

Manuscript draft CLG, DB, LW, AH, JPK, LP, CH, JPK, PL, JHT, ELD

Manuscript revision CLG, DB, RBS, AH, AML, JCYT, LP, MAB, CH, JPK, PL, JHT, ELD

Approve final manuscript – all authors.

All authors take responsibility for their contributions as outlined above.

## Completing Interests

Authors have no competing interest to declare

