## Supplementary material for "A rare *SMAD9* mutation identifies the BMP signalling pathway as a potential osteoanabolic target": All supplemental information

**Supplementary Methods 1**

***High Bone Mass cases from the UK***

DXA databases containing 335,115 DXA scans were initially searched for individuals with a BMD T or Z-score ≥+4 at any site within the lumbar spine (LS) or hip, at 13 centres in England and Wales (9 Hologic, 4 Lunar). A further two centres with Hologic scanners contributed 23 similar individuals identified prospectively. All 1505 DXA images were visually inspected; 962 cases with significant osteoarthritis and/or other causes of raised BMD were excluded (*e.g*. surgical metalwork, Paget's disease, metastases [1]). Evidence of significant osteoarthritis (OA) on lumbar DXA scans is common. To reduce contamination of remaining DXA scans by more moderate OA, we refined case definition by restricting analyses to specific lumbar vertebra(e). At the largest centre, 562 scans with T/Z-score ≥+4 were graded for OA severity and examined in relation to BMD at lumbar vertebral levels [2,3]. In contrast to other lumbar vertebrae, L1 Z-score was not associated with OA, reflecting the recognized pattern of progressive OA changes seen in descending sequential lumbar vertebrae [4]. Further, presence of high TH BMD did not correlate with co-existent LS OA. A standard deviation of +3.2 would be expected to identify a tail of 0.069% of a normal distribution [5]. Of 533 unexplained HBM index cases invited to participate, 240 (45%) were recruited between September 2008 and April 2010 [6]. Written informed consent was collected for all in line with the Declaration of Helsinki [7]. Participants were excluded if under 18 years of age, pregnant or unable to provide written informed consent. All participants were assessed using a standardised structured history and examination questionnaire. DNA was extracted from peripheral venous blood using standard phenol/chloroform extraction techniques.

**Supplementary Methods 2**

**SMAD9_Ex2 Primers**

Fwd primer: CACCCTGTTCAAGGGCTTAG

Rvs primer: CGCGACAGTAAATCACATGG

***PCR Reaction:***

| | |
|---|---|
| $dH_2O$ (PCR grade) | 12.2 |
| 10x ImmoBuffer | 2.5 |
| dNTPs<br>(2.5 mM each dNTP, 10 mM total dNTPs) | 2.5 |
| $MgCl_2$ (50 mM) | 0.75 |
| F (5 μM) | 1 |
| R (5 μM) | 1 |
| Immolase Taq (5 U / μL) | 0.05 |
| DNA template (10 ng /μL) (50ng total) | 5 |
| **Total** | **25** |

***PCR Cycling conditions:***

1. 96˚C for 10 minutes
2. 96˚C for 45 seconds
3. 60˚C for 45 seconds
4. 72˚C for 60 seconds

Go to step 2, 34 more times (35 cycles)

5. 72˚C for 10 minutes
6. 4°C Hold

**Supplementary Results 1: Clinical phenotypes**

***III.1: Index Case (c.65T>C, p.Leu22Pro)***

The 33-year-old index case, with BMD Z-Scores +3.2 at the total hip and +4.5 at L1, had only sustained one traumatic fracture aged 20 months. She reported lower leg and ankle pain. Other than myopia corrected by glasses, she had no visual or auditory impairments, no significant dental history, no back pain or neuropathy, and no pulmonary disease. She had a history of bipolar disorder, polycystic ovary syndrome and psoriasis (no joint involvement), with corresponding medications. She had large feet (shoe size UK 10, Euro 42-43, US 12). She was tall and obese (BMI 43.6), with a broad frame, enlarged mandible and a 4mm torus mandibularis. She had normal joints and no evidence of nerve impingement.

***II.2: Mother of the index case (c.65T>C, p.Leu22Pro)***

The 55-year-old mother, with BMD Z-Scores +3.3 at the total hip and at L1, had never sustained a fracture. Six years earlier she had had a right calcaneal spur surgically removed. She had widespread joint pains affecting ankles, knees (with previous arthroscopy), hips, shoulders, hands and feet, limiting mobility to 50 yards, with a diagnosis of fibromyalgia prompting high analgesic use. She had a history of asthma, hypertension, hypercholesterolaemia and depression with corresponding medications, with normal menopause at 53. She had no visual or auditory impairments, or dental history of note. She was tall and obese (BMI 41.4), with above average shoe size, a broad frame, enlarged mandible but no tori. She had a full range of movement in all joints, bilateral knee crepitus, bilateral pes planus and no signs of nerve compression.

***III.2: Half-sister to index case (c.65T>C, p.Leu22Pro)***

The 22-year-old half-sister, with BMD Z-Scores +4.8 at the total hip and +2.6 at L1, had not fractured. She had had sciatica for five years, lumbar back pain and fronto-temporal headaches for 11 years, with a diagnosis of migraine. She had no significant dental history, no visual or auditory impairments, but was unable to float. She had had a normal menarche at 12. She used inhalers for asthma and medication for depression and anxiety. She was tall and obese (BMI 43), with above average shoe size, a broad frame, enlarged mandible, a torus palitinus in the midline of her hard palate (3cm x 7mm), normal joint movement and no signs of nerve compression.

***I.2: Grandmother of index case (wild-type)***

The 75-years-old grandmother, who did not have HBM (BMD Z-Scores +0.1 at the total hip and +0.8 at L1) had never sustained a fracture. She had osteoarthritis affecting knees, hips, lumbar spine, fingers, left elbow, limiting mobility to 3 metres and a wheelchair. She had had auditory impairment since school and had had cataracts removed. She was overweight (BMI 28.1) with normal shoe size, a normal frame, mandible and no tori, with reduced extension of the right elbow and left knee, with bilateral crepitus of her knees. She died aged 81 of pneumonia secondary to advanced Emphysema.

***Isolated HBM case (c.65T>C, p.Leu22Pro) from the UK***

This 55-year-old female, with BMD Z-Scores +5.0 at the total hip and +4.7 at L1, had never fractured and reported difficulty floating. Her adult left upper cuspid tooth had never erupted; wisdom teeth had been extracted for overcrowding. She had noticed her own mandible enlargement. She had a congenital astigmatism of her left eye with poor vision, and congenital bilateral pes planus. She had no auditory impairment. She took tamoxifen post-surgery for breast cancer. She was obese (BMI 35) with a broad frame, mandible enlargement, but no tori. She had normal joints and no signs of nerve compression.

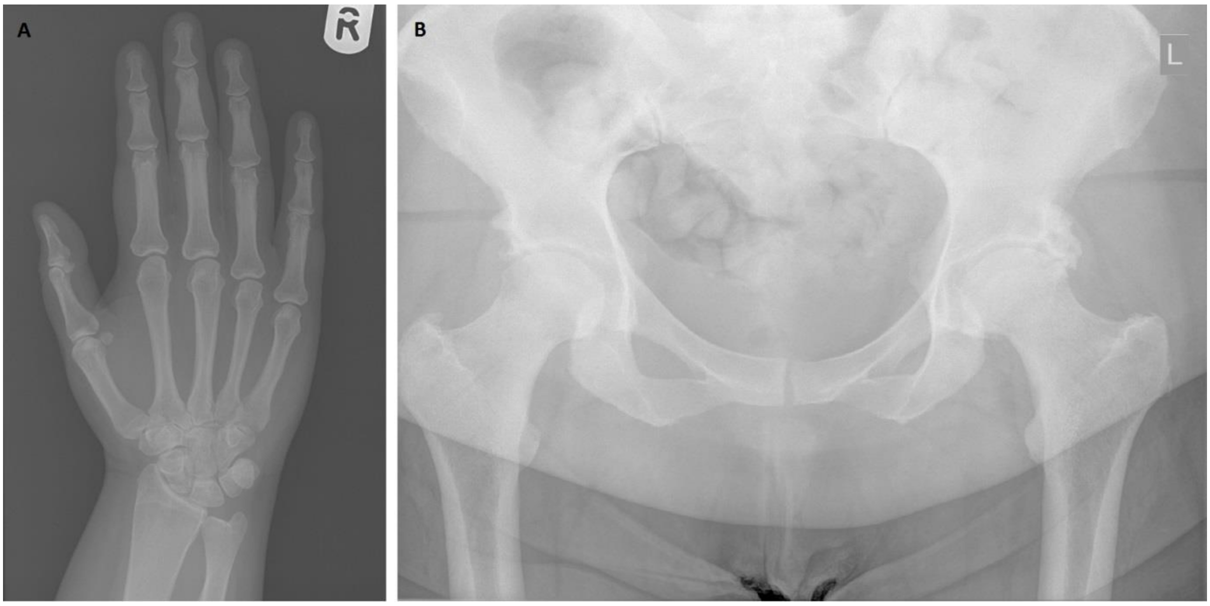


**Supplementary Figure 1 – Plain Radiographs of the (A) right hand and (B) pelvis**

(A) shows prominent metacarpal and proximal phalangeal cortical thicknesses. (B) shows generalised increased density within the iliac blades and thick proximal femoral cortices but no expansive bone changes and normal trabeculation. Bilateral hip joint degenerative changes and new bone formation at the anterior inferior iliac spines bilaterally

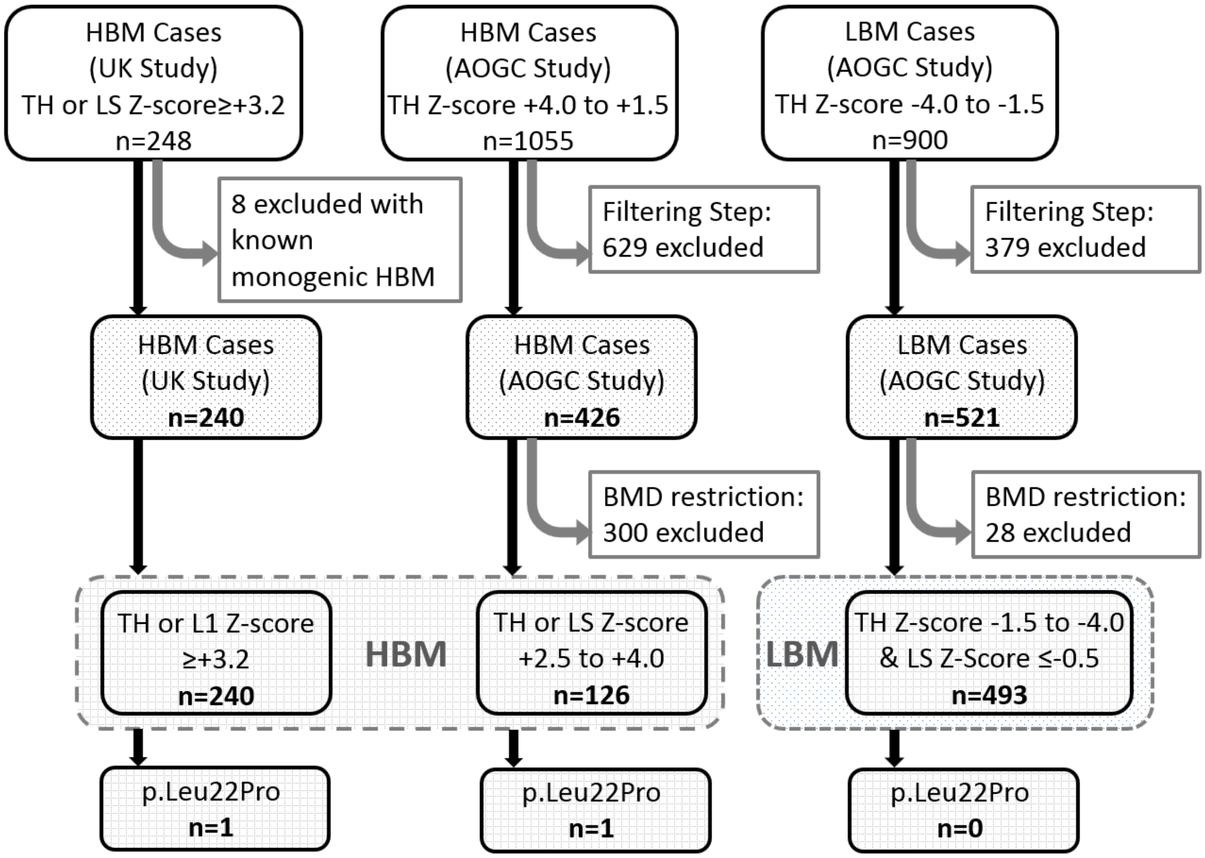


**Supplementary Figure 2 – Flow diagram explaining how further HBM cases were identified for whole-exome sequencing**

TH; Total Hip. LS; Lumbar Spine. QC; Quality Control.
Filtering step restricted sample to individuals who were unrelated, Caucasian ancestry, with complete weight and height data, enough high-quality DNA available for WES, and were able to be sequenced within our financial constraints. BMD restriction ensured all HBM cases had TH or LS Z-score ≥+2.5, and all LBM Cases a TH Z-Score < -1.5 and LS Z-Score ≤-0.5.

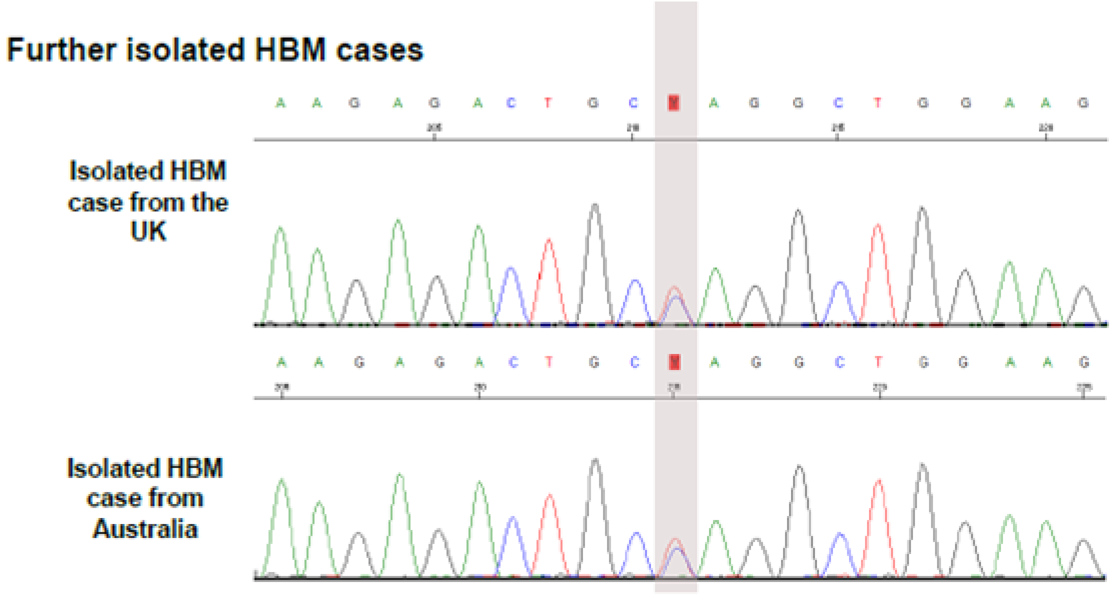


**Supplementary Figure 3 –Electrophoretograms for the additional two isolated unrelated HBM cases with a *SMAD9* c.65T>C, p.Leu22Pro mutation**

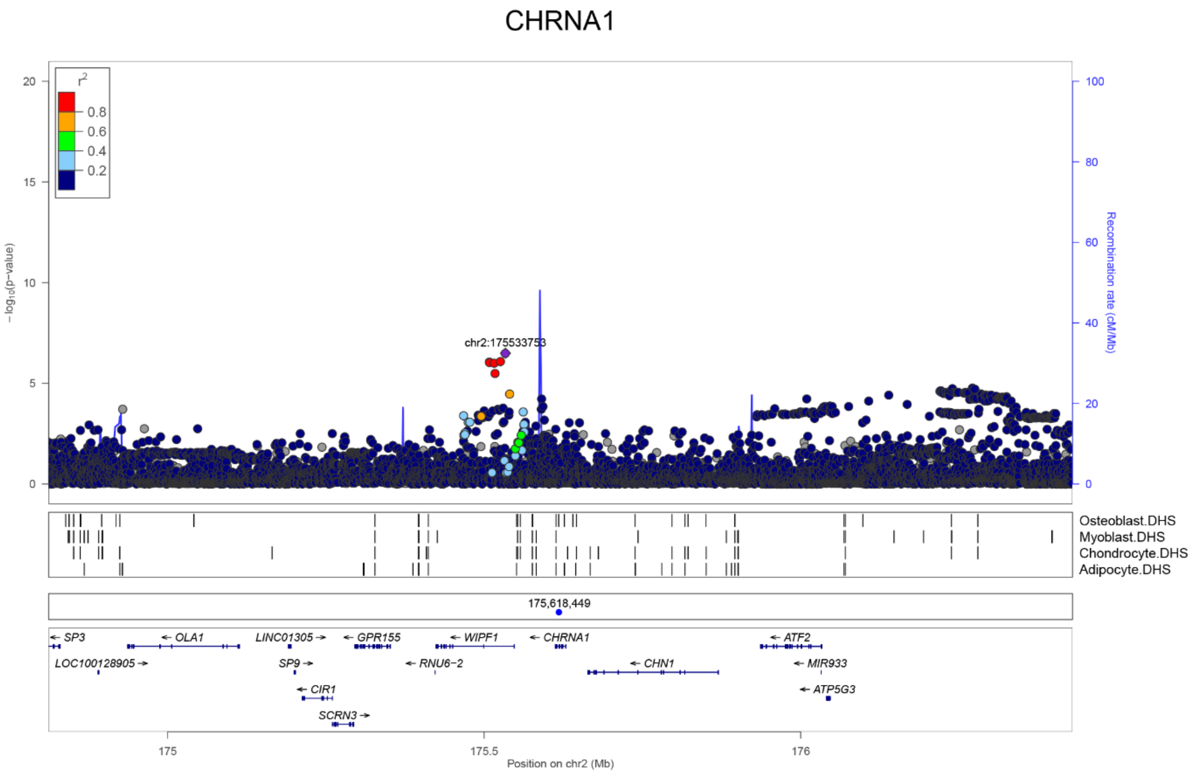


**Supplementary Figure 4 – GWAS for eBMD measured in UK Biobank: Regional association plot for the locus containing *CHRNA1***

Top panel: circles show unconditioned GWAS *P*-values and genomic locations of imputed SNPs within +/- 800kb of the 5' and 3' UTR of each gene. Different colours indicate varying degrees of pairwise linkage disequilibrium between the sentinel eBMD associated SNP (purple diamond) and all other SNPs. Second panel: Vertical shaded areas correspond to locations of DNase I hypersensitive sites (DHSs) characteristic of: skeletal muscle myoblasts cell line (E120), osteoblast primary cells (E129), mesenchymal stem cell derived chondrocyte cultured cells (E049) and mesenchymal stem cell derived adipocyte cultured cells (E023). Red shading depicts intersections between DHSs and genome-wide significant SNPs. Black shading denotes instances in which any other SNPs intersect DHSs. Third panel: Blue circle shows the position of the putative causal mutation c.560T>C, p.Leu187Pro. Fifth panel: Horizontal lines represent genes with vertical lines annotating the location of exons. Arrows indicate the direction in which each gene is transcribed.

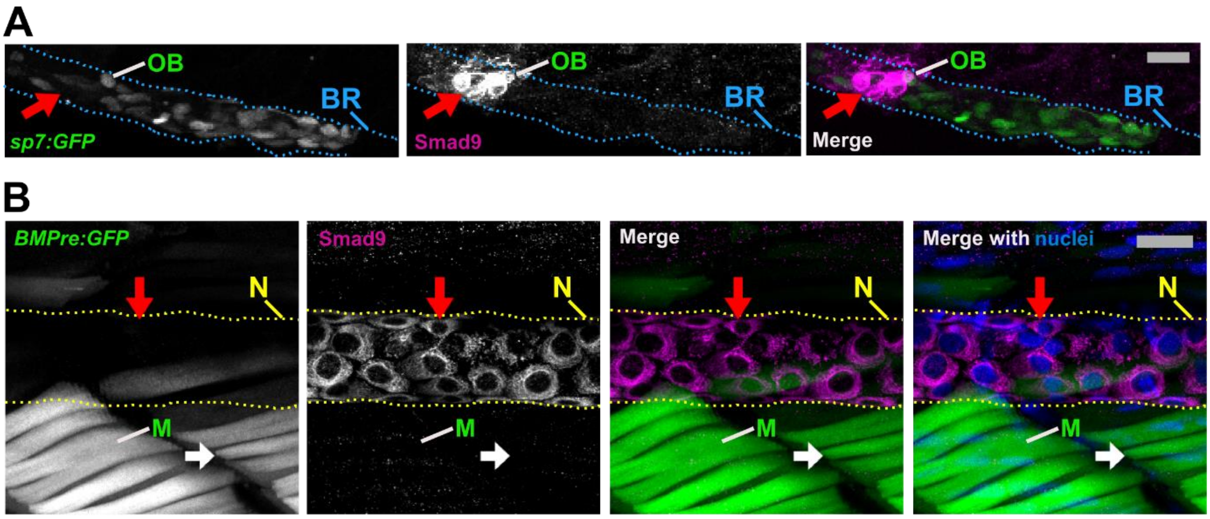


**Supplementary Figure 5 - Smad9 is highly expressed adjacent to osteoblasts in the juvenile branchiostegal ray and in the developing notochordal sheath.**

**A)** High Smad9 expression (magenta, red arrow) in the anterior region of the *sp7:GFP* osteoblast (green) decorated branchiostegal ray (red arrow, intramembranous bone) at 6 days post fertilisation (dpf). Smad9 expression is observed adjacent to *sp7* promoter-driven GFP positive cells (osteoblasts). Blue dotted line outlines the branchiostegal ray.

**B)** At 7 dpf, strong cytoplasmic Smad9 expression was seen in notochordal sheath cells (magenta, red arrow, nucleus DAPI staining in blue) that cover the notochord (yellow dotted line) and which will contribute to vertebrae formation in late larval stages. Smad9-negative and BMP reporter (BMPre) GFP-positive muscle fibres were observed in the trunk muscles adjacent to the cloaca (green, white arrow).

A-B: scale bar is 20 µm, pictures in anterior-posterior (left-right) and dorsal-ventral (top-bottom) orientation from a lateral view, and all maximum intensity z-projection confocal images. Abbreviations: BR, branchiostegal ray; DAPI, 4′,6-diamidino-2-phenylindole; M, muscle; N, notochord; OB, osteoblast.

**Supplementary Video 1 – 3D projection of confocal acquired z-stack showing high Smad9 protein expression adjacent to osteoblasts in the larval zebrafish 6 day old opercle.** Smad9 in magenta and *sp7:GFP* positive osteoblasts in green. 360° rotation rotating at 6° per frame (from Figure 4C image). Note that Smad9 is distinctly separated from the GFP staining.

**Supplementary Video 2: 3-dimensional rotating image of Figure 6**

**Supplementary Table 1: Characteristics measured by pQCT of the p.Leu22Pro *SMAD9* HBM pedigree members and two further isolated p.Leu22Pro *SMAD9* HBM individuals**

| | **HBM Pedigree** | | | | | **Additional Isolated HBM cases** | |
|---|---|---|---|---|---|---|---|
| | UK III.1 | UK III.2 | UK II.2 | UK I.2 | | UK case | Australian case |
| ***SMAD9* Mutation** | Leu22Pro | Leu22Pro | Leu22Pro | WT | | Leu22Pro | Leu22Pro |
| Age | 33 | 22 | 55 | 75 | | 55 | 57 |
| **pQCT 4% Distal Tibia** | | | | | | | |
| Total Bone Area ($mm^2$) | 1051 | 1228 | 1051 | 1118 | | 922 | - |
| Trabecular BMD ($mg/cm^3$) | 361 | 334 | 341 | 269 | | 333 | - |
| Cortical thickness (mm) | 3.29 | 1.90 | 1.59 | 0.02 | | 1.72 | - |
| **pQCT 66% Mid-shaft Tibia** | | | | | | | |
| Total Bone Area ($mm^2$) | 603 | n/a | 610 | 530 | | 612 | - |
| Cortical BMD ($mg/cm^3$) | 1159 | n/a | 1151 | 1025 | | 1139 | - |
| Cortical thickness (mm) | 4.85 | n/a | 4.93 | 2.15 | | 5.10 | - |
| Cortical Bone Area ($mm^2$) | 348 | n/a | 355 | 161 | | 366 | - |
| Cortical/Total Bone Area (%) | 57.7 | n/a | 58.3 | 30.4 | | 59.7 | - |
| SSI ($mm^3$) | 1658 | n/a | 1682 | 756 | | 1700 | - |
| Muscle area ($mm^2$) | 7956 | n/a | 8098 | 4889 | | 8948 | - |

HBM: High Bone Mass, BMD: Bone Mineral Density, SD: Standard Deviation, SSI: Strength Strain Index, WT: Wildtype. n/a, not available as the leg was too large to fit into the pQCT gantry

**Supplementary Table 2: Multi-marker Analysis of GenoMic Annotation (MAGMA) in UK Biobank; summary statistics for all genes within +/- 800kb of *SMAD9***

| GENE ID | ENTREZ ID | CHR | START POSITION | STOP POSITION | No. of SNPs[a] | No. of PARAM[b] | Z STAT[c] | P_SNPWISE_MEAN[d] | P_SNPWISE_TOP1[e] | P_PCREG[f] | P_JOINT[g] |
|---|---|---|---|---|---|---|---|---|---|---|---|
| *SOHLH2* | 54937 | 13 | 36722345 | 36808752 | 636 | 162 | 1.3622 | 2.10E-01 | 4.52E-01 | 2.86E-02 | 8.66E-02 |
| *CCDC169-SOHLH2* | 100526761 | 13 | 36722345 | 36891992 | 1296 | 198 | 1.2181 | 1.13E-01 | 4.96E-01 | 7.71E-02 | 1.12E-01 |
| *CCDC169* | 728591 | 13 | 36781179 | 36891992 | 872 | 142 | 0.8052 | 9.99E-02 | 7.56E-01 | 1.82E-01 | 2.10E-01 |
| *SPG20* | 23111 | 13 | 36855775 | 36964317 | 713 | 175 | 1.9677 | 1.13E-02 | 8.92E-02 | 2.78E-01 | 2.46E-02 |
| *CCNA1* | 8900 | 13 | 36985257 | 37037019 | 321 | 134 | 1.1017 | 9.32E-02 | 9.00E-02 | 6.76E-01 | 1.35E-01 |
| *SERTM1* | 400120 | 13 | 37228049 | 37291976 | 375 | 158 | -0.1693 | 7.64E-01 | 4.81E-01 | 3.27E-01 | 5.67E-01 |
| *RFXAP* | 5994 | 13 | 37373339 | 37423740 | 337 | 134 | -0.36054 | 6.54E-01 | 5.47E-01 | 4.08E-01 | 6.41E-01 |
| ***SMAD9*** | ***4093*** | ***13*** | ***37398968*** | ***37514409*** | ***833*** | ***268*** | ***8.2496*** | ***6.93E-11*** | ***6.65E-11*** | ***1.75E-02*** | ***7.94E-17*** |
| *ALG5* | 29880 | 13 | 37503907 | 37593504 | 533 | 182 | 1.4424 | 4.83E-01 | 7.60E-03 | 5.50E-01 | 7.46E-02 |
| *EXOSC8* | 11340 | 13 | 37554678 | 37603751 | 254 | 116 | 1.8791 | 2.53E-01 | 2.77E-03 | 4.38E-01 | 3.01E-02 |
| *SUPT20H* | 55578 | 13 | 37563449 | 37653850 | 469 | 156 | 1.8396 | 2.10E-01 | 6.98E-03 | 4.07E-01 | 3.29E-02 |
| *CSNK1A1L* | 122011 | 13 | 37657397 | 37699801 | 301 | 88 | -0.021843 | 5.59E-01 | 3.31E-01 | 4.33E-01 | 5.09E-01 |
| *POSTN* | 10631 | 13 | 38116719 | 38192981 | 593 | 151 | 0.39879 | 3.20E-01 | 1.98E-01 | 5.45E-01 | 3.45E-01 |

Using data from 362924 unrelated white British UK Biobank participants.

[a] The number of SNPs annotated to the gene

[b] The number of parameters used in the model

[c] The Z-value for the gene, based on its (permutation) p-value

[d] P-value derived from SNP-wise mean χ2 model

[e] P-value derived from SNP-wise top χ2 model

[f] P-value derived from principal components linear regression model

[g] Aggregate p-value derived from all three methods above (*i.e.* d – f)

**Supplementary Table 3: Multi-marker Analysis of GenoMic Annotation (MAGMA) in UK Biobank; summary statistics for all genes within +/- 800kb of *CHRNA1***

| GENE ID | ENTREZ ID | CHR | START POSITION | STOP POSITION | No. of SNPs[a] | No. of PARAM[b] | Z STAT[c] | P_SNPWISE_MEAN[d] | P_SNPWISE_TOP1[e] | P_PCREG[f] | P_JOINT[g] |
|---|---|---|---|---|---|---|---|---|---|---|---|
| *SP3* | 6670 | 2 | 174751187 | 174850430 | 797 | 207 | 0.7028 | 1.98E-01 | 6.34E-01 | 1.36E-01 | 2.41E-01 |
| *LOC100128905* | 100128905 | 2 | 174870014 | 174911886 | 275 | 111 | 2.6817 | 1.39E-03 | 4.01E-01 | 2.39E-02 | 3.66E-03 |
| *OLA1* | 29789 | 2 | 174917175 | 175133365 | 1375 | 211 | -0.54926 | 5.00E-01 | 6.61E-01 | 5.41E-01 | 7.09E-01 |
| *SP9* | 100131390 | 2 | 175179769 | 175225859 | 305 | 116 | -0.96924 | 9.23E-01 | 6.16E-01 | 5.90E-01 | 8.34E-01 |
| *CIR1* | 9541 | 2 | 175192878 | 175280443 | 519 | 152 | -1.0406 | 8.16E-01 | 7.63E-01 | 5.27E-01 | 8.51E-01 |
| *SCRN3* | 79634 | 2 | 175240457 | 175314303 | 475 | 122 | -0.88671 | 6.44E-01 | 7.47E-01 | 6.32E-01 | 8.12E-01 |
| *GPR155* | 151556 | 2 | 175276299 | 175371816 | 562 | 171 | 0.022405 | 5.05E-01 | 4.71E-01 | 3.31E-01 | 4.91E-01 |
| *WIPF1* | 7456 | 2 | 175404302 | 175567667 | 847 | 255 | 2.5592 | 1.28E-03 | 1.38E-01 | 1.08E-01 | 5.25E-03 |
| ***CHRNA1*** | **1134** | **2** | **175592320** | **175649200** | **380** | **116** | **3.0694** | **3.98E-03** | **6.32E-02** | **4.94E-03** | **1.07E-03** |
| *CHN1* | 1123 | 2 | 175644042 | 175890671 | 1320 | 292 | 1.0285 | 1.35E-01 | 7.73E-01 | 7.36E-02 | 1.52E-01 |
| *ATF2* | 1386 | 2 | 175916978 | 176052934 | 887 | 192 | 0.45519 | 2.47E-01 | 5.37E-01 | 2.29E-01 | 3.24E-01 |
| *ATP5G3* | 518 | 2 | 176020986 | 176066490 | 323 | 104 | -0.61331 | 3.70E-01 | 8.61E-01 | 6.53E-01 | 7.30E-01 |

Using data from 362924 unrelated white British UK Biobank participants.
[a] The number of SNPs annotated to the gene
[b] The number of parameters used in the model
[c] The Z-value for the gene, based on its (permutation) p-value
[d] P-value derived from SNP-wise mean $\chi 2$ model
[e] P-value derived from SNP-wise top $\chi 2$ model
[f] P-value derived from principal components linear regression model
[g] Aggregate p-value derived from all three methods above (*i.e.* d – f)

**Supplementary Table 4: PheWAS for variant rs12427846 using PhenoScanner**

| Trait | EFO | Study | PMID | Ancestry | Year | Beta | SE | P value | Direc-tion | n | N Cases | N Controls | N Studies | Unit | Data set |
|---|---|---|---|---|---|---|---|---|---|---|---|---|---|---|---|
| Forearm bone mineral density | EFO_0007933 | GEFOS | 26367794 | European | 2015 | 0.0648 | 0.0179 | 0.00038 | + | 8143 | 0 | 8143 | 5 | Z-score | A |
| Acute myocardial infarction | EFO_0000612 | Neale B | UKBB | European | 2017 | 0.0012 | 0.0003 | 5.38E-05 | + | 337199 | 3927 | 333272 | 1 | - | B |
| Arm fat-free mass left | - | Neale B | UKBB | European | 2017 | -0.0072 | 0.0018 | 6.54E-05 | - | 331159 | 0 | 331159 | 1 | IVNT | B |
| Arm fat-free mass right | - | Neale B | UKBB | European | 2017 | -0.0059 | 0.0018 | 0.00078 | - | 331221 | 0 | 331221 | 1 | IVNT | B |
| Arm predicted mass left | - | Neale B | UKBB | European | 2017 | -0.0071 | 0.0018 | 7.60E-05 | - | 331146 | 0 | 331146 | 1 | IVNT | B |
| Basal metabolic rate | EFO_0007777 | Neale B | UKBB | European | 2017 | -0.0077 | 0.0019 | 3.77E-05 | - | 331307 | 0 | 331307 | 1 | IVNT | B |
| Chest pain or discomfort | HP_0100749 | Neale B | UKBB | European | 2017 | 0.0035 | 0.0010 | 0.00082 | + | 334053 | 51937 | 282116 | 1 | - | B |
| Heel bone mineral density | EFO_0003923 | Neale B | UKBB | European | 2017 | 0.0148 | 0.0036 | 4.68E-05 | + | 194398 | 0 | 194398 | 1 | IVNT | B |
| Heel bone mineral density left | EFO_0003923 | Neale B | UKBB | European | 2017 | 0.0199 | 0.0048 | 4.05E-05 | + | 106254 | 0 | 106254 | 1 | IVNT | B |
| Height | EFO_0004339 | Neale B | UKBB | European | 2017 | -0.0077 | 0.0020 | 0.00014 | - | 336474 | 0 | 336474 | 1 | IVNT | B |
| Leg fat-free mass left | - | Neale B | UKBB | European | 2017 | -0.0078 | 0.0019 | 3.23E-05 | - | 331258 | 0 | 331258 | 1 | IVNT | B |
| Leg fat-free mass right | - | Neale B | UKBB | European | 2017 | -0.0072 | 0.0019 | 0.00010 | - | 331285 | 0 | 331285 | 1 | IVNT | B |
| Leg predicted mass left | - | Neale B | UKBB | European | 2017 | -0.0077 | 0.0019 | 3.08E-05 | - | 331253 | 0 | 331253 | 1 | IVNT | B |
| Leg predicted mass right | - | Neale B | UKBB | European | 2017 | -0.0072 | 0.0019 | 9.44E-05 | - | 331285 | 0 | 331285 | 1 | IVNT | B |
| Other acute ischaemic heart diseases | EFO_0003777 | Neale B | UKBB | European | 2017 | 0.0003 | 8.3E-05 | 0.00018 | + | 337199 | 288 | 336911 | 1 | - | B |
| Self-reported heart attack or MI | - | Neale B | UKBB | European | 2017 | 0.0016 | 0.0004 | 0.00011 | + | 337159 | 7735 | 329424 | 1 | - | B |
| Treatment with ascorbic acid product | NCIT_C11902; NCIT_C11829 | Neale B | UKBB | European | 2017 | 0.0002 | 4.1E-05 | 0.00025 | + | 337159 | 70 | 337089 | 1 | - | B |
| Treatment with nicorandil | - | Neale B | UKBB | European | 2017 | 0.0006 | 0.0002 | 0.00015 | + | 337159 | 1086 | 336073 | 1 | - | B |
| Trunk fat-free mass | - | Neale B | UKBB | European | 2017 | -0.0073 | 0.0018 | 4.79E-05 | - | 331030 | 0 | 331030 | 1 | IVNT | B |
| Trunk predicted mass | - | Neale B | UKBB | European | 2017 | -0.0072 | 0.0018 | 5.05E-05 | - | 330995 | 0 | 330995 | 1 | IVNT | B |
| Vascular or heart problems diagnosed by doctor: heart attack | - | Neale B | UKBB | European | 2017 | 0.0017 | 0.0004 | 7.08E-05 | + | 336683 | 7790 | 328893 | 1 | - | B |
| Whole body fat-free mass | - | Neale B | UKBB | European | 2017 | -0.0075 | 0.0018 | 2.69E-05 | - | 331291 | 0 | 331291 | 1 | IVNT | B |
| Whole body water mass | - | Neale B | UKBB | European | 2017 | -0.0076 | 0.0018 | 2.08E-05 | - | 331315 | 0 | 331315 | 1 | IVNT | B |
| Coronary artery disease | EFO_0000378; EFO_0001645 | Nelson CP | 28714975 | Mixed | 2017 | 0.0333 | 0.0100 | 0.00088 | + | 148715 | 10801 | 137914 | 1 | log OR | C |
| Coronary artery disease | EFO_0000378; EFO_0001645 | van der Harst P | 29212778 | Mixed | 2018 | 0.03844 | 0.0083 | 3.20E-06 | + | 296525 | 34541 | 261984 | 1 | log OR | D |
| Coronary artery disease | EFO_0000378; EFO_0001645 | van der Harst P | 29212778 | Mixed | 2018 | 0.0319 | 0.0068 | 2.83E-06 | + | 547261 | 122733 | 424528 | 2 | log OR | E |

rsID: rs12427846. hg19_coordinates: chr13:37491650. hg38_coordinates: chr13:36917513. A1: C. A2:T.
Datasets: A: GEFOS_FA_EUR_2015. B: Neale-B_UKBB_EUR_2017. C: Nelson-CP_CAD_Mixed_2017. D: van-der-Harst-P_CAD-UKBB_Mixed_2018. E: van-der-Harst-P_CAD_Mixed_2018

**Supplementary Table 5: - Murine osteocyte expression by whole transcriptome sequencing of Smad9 and Chrna1 in four bone types (tibia, femur, humerus and calvaria)**

| MGI Gene symbol | Chrom | Skeletal GO | Tibia | | Femur | | Humerus | | Calvaria | | Clean Bone | | Bone And Marrow | | Expressed in all bone types | Enriched in osteocyte samples | Osteocyte Signature |
|---|---|---|---|---|---|---|---|---|---|---|---|---|---|---|---|---|---|
| | | | Activity | Mean FPKM | Activity | Mean FPKM | Activity | Mean FPKM | Activity | Mean FPKM | Activity | Mean FPKM | Activity | Mean FPKM | | | |
| **Smad9** | 3 | GO:0060348 GO:0051216 | Active 8/8 | 6.84 | Active 8/8 | 6.13 | Active 8/8 | 6.96 | Active 8/8 | 2.78 | Active 5/5 | 7.19 | Active 1/5 | 0.37 | **Yes** | **Yes** | **Yes** |
| **Chrna1** | 2 | GO:0050881 | Active 4/8 | 0.51 | Active 2/8 | 0.44 | Inactive | 0.25 | Inactive | 0.05 | Inactive | 0.22 | Inactive | 0.05 | **No** | **No** | **No** |

MGI: Mouse Genome Informatics.
Mouse Ensembl Ids: *Smad9* - ENSMUSG00000027796, *Chrna1*- ENSMUSG00000027107
Skeletal GO: Gene Ontology (http://www.ebi.ac.uk)
Activity: Number of replicates of that bone type with FPKM expression values above active gene threshold.
Clean bone: bone with marrow removed to isolate osteocytes
Bone and marrow: bone cleaned of connective tissue and growth plates with the marrow left intact
Mean FPKM: Mean gene expression in for gene across samples of bone type normalised for gene length and library size (Fragments Per Kilobase per Million mapped reads)

**Supplementary Information - Acknowledgements**
We would like to thank all study participants who provided DNA and clinical information. Regarding the HBM study, we particularly thank staff at the Wellcome Trust Clinical Research Facility in Birmingham, Royal National Hospital for Rheumatic Diseases in Bath, Cambridge NIHR Biomedical Research Centre and Addenbrooke's Wellcome Trust Clinical Research Facility in Cambridge, Bone Research Unit in Cardiff, Musculoskeletal Research Unit in Bristol, NIHR Bone Biomedical Research Unit in Sheffield and the Brocklehurst Centre for Metabolic Bone Disease in Hull.

Regarding the Anglo-Australasian Genetics Consortium (AOGC) study, we thank the AOGC PIs; Eugene McCloskey (Sheffield, UK), Geoffrey C Nicholson (Geelong, Australia), Richard Eastell (Sheffield, UK), Richard L Prince (Perth, Australia), John A Eisman (Sydney, Australia), Graeme Jones (Hobart, Australia), Philip Sambrook (Sydney, Australia), Ian R Reid (Auckland, New Zealand), Elaine M Dennison (Southampton, UK), John Wark (Geelong, Australia). Furthermore, we thank Barbara Mason and Amanda Horne (Auckland) for patient recruitment; Judith Finigan (Sheffield, UK) for laboratory support and database support; Selina Simpson (Sheffield, UK) for DNA handling; Fatma Gossiel (Sheffield, UK) for DNA handling; Alison Steward and Lana Gibson (Aberdeen, UK) for patient recruitment; Katherine Kolk (Geelong, Australia); Janelle Rampellini (Perth, Australia) for patient recruitment; Jemma Christie (Melbourne, Australia) for patient recruitment; Helen Steane (Hobart, Australia) for patient recruitment; Denia Mang and Ruth Toppler for DNA extraction, DNA handling, and database support (Dubbo/Sydney, Australia); Kate Lowings (Brisbane, Australia) for patient recruitment; and Marieke Brugmans and Leanne Brookes (Brisbane, Australia) for DNA preparation and genotyping. We thank Ms Linda Bradbury (Brisbane, Australia) for support with ethics, governance and recruitment.

The HBM study was supported by the UK NIHR CRN (portfolio number 5163); supporting CLRNs included Birmingham and the Black Country, London South, Norfolk & Suffolk, North and East Yorkshire and Northern Lincolnshire, South Yorkshire, Surrey & Sussex, West Anglia and Western.

The AOGC also received funding from the Australian Cancer Research Foundation and Rebecca Cooper Foundation (Australia). MAB was funded by a National Health and Medical Research Council (Australia) Principal Research Fellowship and ELD was funded by a National Health and Medical Research Council (Australia) Career Development Award (569807). IR is supported by the Health Research Council of New Zealand. The OPUS study was supported by Sanofi-Aventis, Eli Lilly, Novartis, Pfizer, Proctor & Gamble Pharmaceuticals and Roche. The Sydney Twin Study was supported by the National Health and Medical Research Council, Australia. The Dubbo Osteoporosis Epidemiology Study was supported by the Australian National Health and Medical Research Council, MBF Living Well foundation, the Ernst Heine Family Foundation and from untied educational grants from Amgen, Eli Lilly International, GE-Lunar, Merck Australia, Novartis, Sanofi-Aventis Australia and Servier. The Hertfordshire Cohort Study was supported by grants from the Medical Research Council UK & Arthritis Research UK. The Geelong Osteoporosis Study was funded by grants from the Victorian Health Promotion Foundation and the Geelong Region Medical Research Foundation, and the National Health and Medical Research Council, Australia (project grant 628582). The Oxford Osteoporosis Study was funded by Action Research UK.

This research has been conducted using the UK Biobank Resource (accession IDs: 12703). We would like to thank the Wolfson Bioimaging Facility at the University of Bristol, UK, for confocal microscope access and imaging support and Jessica Harris and Sharon Song at the Queensland University of Technology for their help with genotyping.

**Supplementary Information References**

1 Gregson, C. L., Hardcastle, S. A., Cooper, C. & Tobias, J. H. Friend or foe: high bone mineral density on routine bone density scanning, a review of causes and management. *Rheumatology* **52**, 968-985 (2013).

2 Kellgren, J. & Lawrence, J. Radiological assessment of osteo-arthrosis. *Ann Rheum Dis* **16**, 494-502 (1957).

3 Gregson, C. L., Steel, S., Yoshida, K., Reid, D. M. & Tobias, J. H. An investigation into the impact of osteoarthritic changes on bone mineral density measurements in patients with High Bone Mass. *ASBMR 30th Annual Meeting, Montreal.* **SA257** (2008).

4 Hansen, K. E. *et al.* Interobserver reproducibility of criteria for vertebral body exclusion. *J Bone Miner Res.* **20**, 501-508 (2005).

5 White, J., Yeats, A. & Skipworth, G. *Tables for Statisticians*. Vol. 3rd (Stanley Thornes, Cheltenham, 1979).

6 Gregson, C. L. *et al.* 'Sink or swim': an evaluation of the clinical characteristics of individuals with high bone mass. *Osteo Int.* **23**, 643-654 (2012).

7 The 59th General Assembly Seoul. World Medical Assembly Declaration of Helsinki - Ethical Principles for Medical Research Involving Human Subjects., 2008).
